# GABAergic neurons and Na_V_1.1 channel hyperactivity: a novel neocortex-specific mechanism of Cortical Spreading Depression

**DOI:** 10.1101/2020.03.14.991158

**Authors:** Oana Chever, Sarah Zerimech, Paolo Scalmani, Louisiane Lemaire, Alexandre Loucif, Marion Ayrault, Martin Krupa, Mathieu Desroches, Fabrice Duprat, Sandrine Cestèle, Massimo Mantegazza

## Abstract

Cortical spreading depression (CSD) is a pathologic mechanism of migraine. We have identified a novel neocortex-specific mechanism of CSD initiation and a novel pathological role of GABAergic neurons. Mutations of the Na_V_1.1 sodium channel (the *SCN1A* gene), which is particularly important for GABAergic neurons’ excitability, cause Familial Hemiplegic Migraine type-3 (FHM3), a subtype of migraine with aura. They induce gain-of-function of Na_V_1.1 and hyperexcitability of GABAergic interneurons in culture. However, the mechanism linking these dysfunctions to CSD and FHM3 has not been elucidated. Here, we show that Na_V_1.1 gain-of-function, induced by the specific activator Hm1a, or mimicked by optogenetic-induced hyperactivity of cortical GABAergic neurons, is sufficient to ignite CSD by spiking-generated extracellular K^+^ build-up. This mechanism is neocortex specific because, with these approaches, CSD was not generated in other brain areas. GABAergic and glutamatergic synaptic transmission is not required for optogenetic CSD initiation, but glutamatergic transmission is implicated in CSD propagation. Thus, our results reveal the key role of hyper-activation of Na_v_1.1 and GABAergic neurons in a novel mechanism of CSD initiation, which is relevant for FHM3 and possibly also for other types of migraine.

## Introduction

Cortical spreading depression (CSD) is a wave of transient intense network hyperexcitability leading to a long lasting depolarization block of neuronal firing, which initiates focally and then slowly propagates in the cerebral cortex (Pietrobon and Moskowitz, 2014). CSD is the cause of migraine aura and could induce migraine headache by sensitization of meningeal nociceptors (Ferrari et al., 2015; Pietrobon and Moskowitz, 2014). CSD events, as well as spreading depolarizations generated in anoxic conditions, have been observed and extensively studied for decades, but specific pathological mechanisms that lead to their initiation are still far from being completely understood and sorted out (Dreier and Reiffurth, 2015). A mendelian form of migraine with aura (MA) characterized by hemiparesis during the attacks, Familial Hemiplegic Migraine (FHM), has allowed the identification of some molecular/cellular pathological mechanisms (Ferrari et al., 2015; Pietrobon and Moskowitz, 2014), and has become a model disease for more common forms and for studies of mechanisms of CSD. FHM type 1 (FHM1) is caused by gain of function mutations of the α1 subunit of the Ca_V_2.1 P/Q type Ca^2+^ channel (the *CACNA1A* gene) (Ophoff et al., 1996), FHM type 2 (FHM2) by loss-of-function mutations of the α2 subunit of the glial Na^+^/K^+^ pump (the *ATP1A2* gene) (De Fusco et al., 2003). The experimental induction of CSD is facilitated in both FHM1 and FHM2 genetic mouse models (Leo et al., 2011; van den Maagdenberg et al., 2004), primarily because of increased network excitability which is induced by excessive release or insufficient reuptake of glutamate (Capuani et al., 2016; Tottene et al., 2009; Vecchia et al., 2014), consistent with a similar overall mechanism affecting the glutamatergic system. FHM3 is caused by mutations of Na_V_1.1 (*SCN1A*) Na^+^ channel (Dichgans et al., 2005), which is particularly important for GABAergic neurons’ excitability (Yu et al., 2006). Numerous epileptogenic Na_V_1.1 mutations have been identified, and studies performed both *in vitro* and in mouse models have shown that they cause loss-of-function of the channel, leading to decreased excitability of GABAergic neurons, reduced inhibition and consequent hyperexcitability of cortical networks (Catterall et al., 2010; Hedrich et al., 2014; Mantegazza, 2011; Mantegazza and Broccoli, 2019; Ogiwara et al., 2007; Yu et al., 2006). We and others have provided evidence that, in contrast to epileptogenic mutations, FHM3 mutations cause instead gain-of-function of the channel, often increasing persistent current, and hyperexcitability of transfected GABAergic neurons in primary culture (Barbieri et al., 2019; Bertelli et al., 2019; Cestele et al., 2013a; Cestele et al., 2008; Cestele et al., 2013b; Dhifallah et al., 2018; Fan et al., 2016; Mantegazza and Broccoli, 2019; Mantegazza and Cestele, 2017), which could be responsible of CSD initiation, as we have recently proposed in a computational model (Desroches et al., 2019). Interestingly, a recent work shows that knock-in mice carrying the FHM3 L263V Na_V_1.1 mutation experience spontaneous CSD events but not seizures, although detailed mechanisms of CSD generation have not been studied (Jansen et al., 2019). Thus, these results point to different, novel and counterintuitive mechanisms in comparison to FHM1 and FHM2 mutations. However, there is not yet a causal link between Na_V_1.1 gain-of-function/GABAergic neurons’ hyperactivity and CSD, and it is not clear whether and how CSD could be generated by these dysfunctions. In particular, this would be a novel role of GABAergic neurons, hitherto unknown, implicated in the generation of CSD. Here, we addressed these issues inducing acute Na_V_1.1 gain-of-function and performing selective optogenetic stimulation of GABAergic neurons, carrying out experiments *in vivo*, *ex vivo* in brain slices and *in vitro* in cell lines.

## Materials and Methods

### Animal care and mouse lines

Experiments were carried out according to the European directive 2010/63/UE and approved by institutional and ethical committees (approval #C06-152-5 for France, 711/2016-PR for Italy). All efforts were made to minimize the number of animals used and their suffering. They were group housed (5 mice per cage, or 1 male and 2 females per cage for breeding) on a 12 h light/dark cycle, with water and food *ad libidum*. Mouse lines have been obtained from Jackson laboratory (JAX, USA), besides GAD67-GFP knock-in mice (Tamamaki et al., 2003), which have been obtained from Yuchio Yanagawa (Gunma University, Japan).

To have specific expression of channelrhodopsin-H134R/tdtomato in GABAergic neurons, most of the experiments have been performed with male and female double hemizygous transgenic mice VGAT-hChR2(H134R)/tdtomato (VGAT-ChR2 in the text) and control littermates (F1 generation). They were obtained by mating hemizygous females loxP-STOP-loxP-hChR2(H134R)-tdtomato (Ai27D, B6.Cg-Gt(ROSA)26Sortm27.1(CAG-COP4*H134R/tdTomato)Hze/J, JAX n°012567) (Madisen et al., 2012) with transgenic males (to avoid off-target Cre expression in the female germline) hemizygous for the Viaat-Cre transgene (Cre recombinase expression driven by the vesicular GABA transporter, VGAT, promoter) (B6.FVB-Tg(Slc32a1-Cre)2.1Hzo/FrkJ, JAX n°017535), line 2.1 in (Chao et al., 2010). To avoid germline transmission of recombined floxed alleles (see www.jax.org/strain/017535), we have never used VGAT-ChR2 F2 offspring. VGAT-Cre mice have been previously used for obtaining specific expression of floxed alleles in GABAergic neurons (Baalman et al., 2015; Chao et al., 2010; Li et al., 2016; Wagnon et al., 2011; Wang et al., 2014). To evaluate the effect of a reduction of Na_V_1.1 expression in interneurons, we crossed double hemizygous VGAT-ChR2 mice with heterozygous Na_V_1.1 knock-out mice (*Scn1a*^+/−^) (Yu et al., 2006). Moreover, for immunohistochemistry, we used mice expressing a floxed td-tomato transgene (Ai9, B6;129S6-Gt(ROSA)26Sor^tm9(CAG-tdTomato)Hze^/J; Jackson Lab. n°007905) as a cell filling reporter line, because it was difficult to identify hChR2(H134R)/tdtomato expressing cells with the plasma membrane fluorescence of the tagged ChR2 in VGAT-ChR2 mice. All mouse lines were in the C57BL/6J background (>10 generations, Charles River, USA), besides *Scn1a*^+/−^ mice that were in a mixed background (C57BL/6J-CD1 85:15%).

Offspring was genotyped either by PCR, following the standard JAX protocols, using our standard protocol for *Scn1a*^+/−^ knock-out mice (Liautard et al., 2013) or, for mice with cell filling fluorescent protein, controlling the fluorescence of newborn mice with a Dual Fluorescent Protein Flashlight (NightSea, USA). We used mice of both sexes, 4-6 weeks old for *ex-vivo* experiments and 4-8 weeks old for *in vivo* experiments.

### Preparation of brain slices, electrophysiological recordings and imaging in slices

Brain slices were prepared as previously described (Hedrich et al., 2014; Zerimech et al., 2020). Briefly, mice were killed by decapitation under isoflurane anesthesia, the brain was quickly removed and placed in ice-cold artificial cerebrospinal fluid (ACSF), which contained (in mM): 125 NaCl, 2.5 KCl, 2 CaCl_2_, 1 MgCl_2_, 1.25 NaH_2_PO_4_, 25 NaHCO_3_ and 25 glucose, saturated with 95 % O_2_ −5 % CO_2_. Acute coronal slices 400 μm thick (300 μm for spatial illumination and patch-clamp experiments) were prepared with a vibratome (HM650V, MicroM, Germany) in ice-cold ACSF. Slices were then stored in a submerged chamber with ACSF at 34°C at least one hour before the beginning of the recordings. Selective brain expression of the ChR2(H134R)/tdTomato transgene and thus absence of germline recombination of the floxed allele was systematically confirmed by visual inspection of slices and of other tissues of the mouse using the NightSea Flashlight (USA). Inductions of CSD started > 10 min after placing an individual slice in the recording chamber (RC-26GLP; Warner Instruments, USA), which was perfused with recording ACSF (rACSF) (Tottene et al., 2009) at 34 °C, containing (in mM): 125 NaCl, 3.5 KCl, 1 CaCl_2_, 0.5 MgCl_2_, 1.25 NaH_2_PO_4_, 25 NaHCO_3_ and 25 glucose, saturated with 95 % O_2_ −5 % CO_2_. Slices were visualized using infra-red differential interference contrast (DIC) microscopy (Nikon Eclipse FN1, Japan) equipped with a CCD camera (CoolSnap ES2, Photometrics, USA), filter cubes for visualization of fluorescent proteins (Semrock, USA) and a filter for optogenetic 475 nm blue light illumination (FF02-475/50; Semrock, USA). Acquisitions of large fields of view were obtained with a 0.35X camera adapter lens. Electrophysiological signals were recorded with a Multiclamp 700B amplifier (CV-7B headstage), a Digidata 1440A acquisition board and pClamp 10.3 software (Molecular Devices, USA). DC extracellular field potential recordings were performed using borosilicate glass micropipettes (∼0.5 MΩ) filled with rACSF. Whole-cell patch-clamp recordings were performed using borosilicate glass pipettes of 3–5 MΩ resistance containing (in mM): 120 K-gluconate, 15 KCl, 2 MgCl_2_, 0.2 EGTA, 10 HEPES, 20 P-Creatine, Na_2_-0.2 GTP, 2 Na_2_-ATP, 0.1 leupeptin, adjusted to pH 7.25 with KOH. Whole-cell access resistance (10-25 MΩ) was monitored and cells showing instable access resistance (> 20 %) were discarded. Recordings were started 5 min after obtaining the whole-cell configuration. Ion sensitive electrodes were built with glass pipettes (tip diameter ∼ 2 μm) as previously described (Chever et al., 2010). Briefly, they were first cleaned with absolute ethanol, then pre-treated with dimethylchlorosilane vapors (Sigma-Aldrich, USA), dried at 100 °C for 2 h, and finally the tip was backfilled with the K^+^ ionophore I-cocktail B (Fluka, Sigma-Aldrich, USA) using a 28G microfil (WPI, USA). The calibration was done for each microelectrode using solutions of different K^+^ concentrations (2.5, 3.5, 35, and 90 mM: KCl added to the ACSF), and electrodes were selected according to the slope of their response (a minimum of 50 mV/decade increase in K^+^ concentration). To minimize the leak current of the headstage in ion sensitive recordings, we used Rf = 5 GΩ and compensated the residual leak current: we zeroed the pipette offset observed with a 10 MΩ load, we then switched to a 10 GΩ load and we re-zeroed the offset observed in this condition (which is mainly due to the residual leak current) with the current injection circuitry of the amplifier (holding current); the estimated leak current (equal and opposite to the holding current applied) was 0.73 pA at 25 °C and stable over time. For accurate evaluation of [K^+^]_out_ dynamics, the field potential signal was subtracted from the K^+^-sensitive recording (the ion sensitive and the extracellular potential pipettes were placed at < 100 µm of distance from each other). Electrophysiological signals were filtered at 10 kHz and sampled at 25 kHz. Multi-unit activities (MUA) were obtained band-pass filtering off-line the LFP trace at 300-500 Hz with pClamp (single pole RC filters). Intrinsic optical signal (IOS: near-infrared light transmittance) (Holthoff and Witte, 1996) was monitored acquiring images with the CoolSnap ES2 CCD camera controlled with micromanager (Edelstein et al., 2014) at 1 image/s (unless otherwise indicated). Analyses and signal/image processing were performed using pClamp 10.3, Origin 8 (Origin Lab, USA) and ImageJ-Fiji (Schindelin et al., 2015).

### Optogenetic illumination of brain slices

Activation of ChR2 was obtained illuminating brain slices through the 4x objective. A white light source (130 W mercury lamp, Intensilight, Nikon, Japan) was connected to the epi-illumination port of the microscope with a light guide containing a 420 nm UV blocker filter (series 2000, Lumatec, Germany); the white light was filtered with a 475/50 filter (Semrock) placed in the optical path and delivered to the objective with a FF685-Di02 dichroic beamsplitter (Semrock). The area of illumination was 38.5 mm^2^. The blue light power density measured with a power meter (Ophir Photonics, USA) was 2.8 mW/mm^2^ at the slice surface (photodetector placed at the level of the slice). For spatial illumination experiments, the light guide was connected to a digital micromirror device (DMD)-based patterned photostimulator (Polygon 400, Mightex, Canada), which was connected to the microscope with a custom adapter and a NI-FLT6 Epi-fluorescence Cube Turret equipped with a FF685-Di02 dichroic beamsplitter and a 475/50 filter. The area of the spatial illumination was between 0.34 and 1.1 mm^2^. ON/OFF of the illumination was controlled with the Intensilight shutter or with the Polygon 400 software. We mainly used continuous illumination, but trains of illumination have been also successfully tested (with spatial illumination: 5 Hz, duty cycle 50 %). The infrared filter of the microscope, besides for patch-clamp experiments, was used in most experiments to eliminate the blue light of the optogenetic illumination from the acquired images and to obtain near-infrared IOS.

### Induction of CSD by application of KCl

Cortical spreading depression was induced by brief puffs of KCl (130 mM) in the superficial cortical layers (layers 1-3) with a glass micropipette (2-4 MΩ) connected to an air pressure injector (holding pressure: < 1 PSI, injection pressure: 7 to 10 PSI; PV820 Pneumatic Picopump, WPI, USA) (Zerimech et al., 2020). The field potential recording pipette was placed at least 500 μm away from the CSD induction area. For CSD induction with a solution containing 12 mM KCl (12 mM KCl, 125 mM NaCl, negative control solution was 137 mM NaCl saline solution), long local perfusions of KCl were performed with a glass pipette of 0.5 MΩ (3 PSI) until CSD induction. For both brief puffs and long local perfusions, Fastgreen (0.1 %, SIGMA-Aldrich, USA) was added to the pipette solutions to visualize the injection area. Application of the control NaCl solutions with Fastgreen never ignited CSD.

### Image processing and analyses

Image analysis with ImageJ-Fiji was used for identifying the CSD wave by intrinsic optical imaging. Image processing was performed to evaluate latency of CSD initiation and to quantify CSD propagation speed with a custom made macro (available at https://www.ipmc.cnrs.fr/~duprat/scripts/imagej.htm): to eliminate the background and isolate the wave from the raw image, a representative image acquired before the 470 nm illumination was subtracted from the others and then contrast was enhanced. To determine the speed of the propagating wave, successive line plots of the wave front from processed images (every 2 s) were drawn manually and the spatial distances between them were measured by means of the peak finder ImageJ plugin. For each slice, speed was estimated on a minimum of 4 time points (8 s). For few experiments, the quality of the images was not sufficient for a reliable quantification of CSD propagation speed, and they were considered as “non-available” (NA) data (reported in the supplementary table). Latencies of CSD initiation were quantified off-line using processed images. Only CSDs that initiated in the visual field of the camera were used for latency analysis; we considered the others as “non-available” data (see the supplementary table).

For the evaluation of the percentage of CSD induction, individual slices were exposed to 470 nm illumination for 90 s maximum. When the illumination did not induce CSD within this time limit, we tested that the slice could generate CSD by injecting a puff of 130 mM KCl solution as described above. The percentage of optogenetically induced CSD was quantified considering as non-successful optogenetic inductions only experiments in which the slice could generate CSD with a subsequent application of KCl. The same protocol was used to determine the success rate of spontaneous CSD induction following bath application of Hm1a toxin: individual slices from VGAT-ChR2 mice were perfused with rACSF or with Hm1a (10 nM) dissolved in rACSF for 10-15 min. If no CSD occurred in this time window, then optogenetic CSD induction was tested. The percentage of CSD triggered following Hm1a was quantified considering as non-successful inductions experiments in which the slice could generate CSD with a subsequent 470 nm illumination. “Aborted CSD” refers to CSD that rapidly decelerate after initiation and stop propagating within <800 µm from the initiation site.

### Immunohistochemistry, confocal acquisitions, and cell count

Mice were anesthetized (disodium salt pentobarbital, 40 mg/kg, IP) and intracardially perfused with cold PBS (Sigma-Aldrich, USA) and paraformaldehyde 4 % (Electron Microscopy Sciences, USA) prepared in PBS. Brains were removed and post-fixed overnight. 40 µm-coronal sections were performed with a microtome (HM650V, Microm, Germany). Before immunohistochemistry processing, floating sections were permeabilized and immunoblocked with Triton X-100 (0.1 %, Sigma-Aldrich, USA), NGS (10 %, Normal Goat serum, S-1000, Vector lab, USA), BSA (0.5 %, Bovine serum albumin, Amresco, USA) diluted in PBS, at RT for 30 min. Then, slices were incubated overnight at 4 °C with primary antibody, and after several washes at RT, with the appropriated secondary antibody. Antibodies were diluted in PBS containing Triton X-100 (0.1 %), NGS (1 %), BSA (0.5 %). Slices were mounted on glass slides using Fluoroprep (Biomérieux, France) or Mowiol (Sigma Aldrich, USA) as mounting media. All antibodies used in the study are commercially available and have been validated in previous published studies, as reported by the suppliers.

Primary antibodies used: anti-NeuN (1/1000) mouse monoclonal antibody (MAB377, Merck Millipore, USA), anti-CamKII (1/100) mouse monoclonal antibody (6G9 clone, MA1-048, ThermoFischer Scientific, USA), anti-S100 (1/1000) rabbit polyclonal antibody (Z0311, Dako Corp., USA). Fluorescent dye-conjugated secondary antibodies used in appropriate combinations: Goat-anti mouse or anti-rabbit antibodies (1/1000: Alexa 647, A21236 and A21245, Invitrogen, USA).

Immunofluorescence acquisitions were performed using a confocal laser-scanning microscope FV10i (Olympus, Japan), with a 60x objective (image format: 1024×1024). Mosaics of ∼1.5 mm × 1.5 mm were performed with 635 nm and 559 nm laser lines. Quantification of fluorescent cells and co-localisation of fluorescent markers were performed with ImageJ-Fiji and the cell counter plugin, using 200 µm-width regions of interest (ROI) drawn perpendicularly to the cortical surface and covering all the cortical layers. 4-5 ROI were used for each mouse and the grand average was then performed with 3 mice.

### Patch-clamp recordings in cell-lines

#### Plasmids, cell culture and transfections

The cDNAs of the Na_V_1.1 Na^+^ channel (GenBank sequence NM_006920.4) and of the Na_V_1.2 Na^+^ channel (GenBank sequence NM_021007) were obtained from Jeff Clare (Glaxo-SmithKline, UK) and subcloned into the pCDM8 vector to reduce rearrangements, as already described (Bechi et al., 2012; Cestele et al., 2013b). The plasmids were propagated in MC1061/P3 *E. Coli* (Invitrogen-ThermoFisher, USA) grown at 28 °C for > 48 h, and the entire coding sequence was sequenced after each propagation to rule out the presence of unwanted spurious mutations or rearrangements. Na_V_1.1 or Na_V_1.2 were co-expressed with a reporter vector expressing YFG (pEYFP-N1) in tsA-201 cells (authenticated cells purchased from ECACC-Sigma: cell line 96121229) transiently transfected by using the CaPO_4_ method, as previously described (Bechi et al., 2012; Cestele et al., 2013b); tsA-201 cells were cultured in modified Dulbecco’s medium and Hams-F12 mix supplemented with 10 % fetal bovine serum (Invitrogen-ThermoFisher, USA). Na_v_1.6 Na^+^ channel (GenBank sequence NM_014191.2) was stably expressed in HEK293 cells (Bechi et al., 2012; Oliveira et al., 2004) (obtained under a MTA from Jeff Clare, Glaxo-SmithKline, UK), which were cultured in modified Dulbecco’s medium supplemented with 10 % foetal bovine serum (Invitrogen-ThermoFisher, USA). Cell cultures were tested for mycoplasma contamination (LookOut kit, Sigma-Aldrich).

#### Whole-cell recordings and analysis

Sodium currents were recorded with the whole-cell configuration of the patch-clamp technique (24-48 h after transfection for Na_V_1.1 and Na_V_1.2) as previously described (Bechi et al., 2012; Cestele et al., 2013b). The recordings were performed at room temperature (22-25 °C) using a Multiclamp 700A amplifier and pClamp 10.2 software (Axon Instruments/Molecular Devices). Signals were filtered at 10 kHz and sampled at 100 kHz. Electrode capacitance and series resistance were carefully compensated throughout the experiment. Pipette resistance was between 1.5 and 2.5 MΩ; maximum accepted voltage-clamp error was 2.5 mV. The remaining transient and leakage currents were eliminated online using a P/4 subtraction paradigm. The extracellular recording solution contained (in mM): 150 NaCl, 10 HEPES, 2 KCl, 1.5 CaCl_2_ and 1 MgCl_2_ (pH 7.4 with NaOH); the internal pipette solution contained (in mM): 105 CsF, 35 NaCl, 10 EGTA, 10 HEPES (pH 7.4 with CsOH).

### Pharmacological agents and chemicals

Gabazine, Isoguvacine and CdCl_2_ were bought from Sigma-Aldrich (USA), CNQX disodium salt, VU0240551 and VU0463271 from Tocris Bioscience (UK), CPP and TTX-citrate from Alomone labs (Israel). Synthetic Hm1a was purchased from Smartox S.A.S (France). All other chemicals were purchased from Sigma-Aldrich. Drugs were applied for 15 min before to test the induction of CSD.

### *In vivo* experiments

Mice were deeply anesthetized with ketamine/xylazine (100 mg/kg and 5 mg/kg, respectively) and placed in a Faraday cage-shielded stereotaxic frame. Supplemental doses of anesthesia were applied on appearance of withdrawal reflex in response to limb pinching. The body temperature was maintained at 37 °C with a heating pad; the stability of the respiratory activity was monitored with a piezoelectric transducer (MLT1010 Pulse Transducer, AD Instruments, UK) and that of the heart rate with ECG recordings. A craniotomy was performed (2 mm antero-posterior, 3.5 mm lateral with respect to bregma), the dura was carefully removed, and mineral oil (Sigma-Aldrich) was applied on the cortex to prevent cortical surface drying. A glass pipette filled with 0.9 % NaCl (1-2 µm tip diameter, Ag/AgCl electrode) was lowed into the barrel cortex for DC field potential recordings. The placement of the pipette in the barrel cortex was confirmed recording the spiking activity induced by vibrissae stimulations. A 400 µm diameter optical fiber, connected to a 470 nm LED light source (Optoflash, Cairn research, UK; 30 mW/mm^2^ at the optical fiber tip) was placed on the surface of the cortex, in the area of the recording pipette. The photostimulations consisted of 100 Hz trains of 0.8 ms pulses, delivered till a CSD was triggered; after 100 s the stimulation was considered unsuccessful. Note that trains of illumination were able to induce CSD also in brain slices (supplementary video 4), DC field potentials were recorded with a differential extracellular amplifier (EX4-400, Dagan, USA) and acquired with a Digidata 1440A and pClamp software (Molecular Devices, USA). Anesthetized mice were sacrificed at the end of the recordings by cervical dislocation.

### Computational model

The model of two coupled neurons, GABAergic and pyramidal, was based on the one we developed in (Desroches et al., 2019). Here, we refined the model introducing the modifications described below, and performed the numerical simulations with the software XPPAUT (http://www.math.pitt.edu/~bard/xpp/xpp.html).

In the revised model, we better modeled the dynamics of ionic concentrations, which is an essential feature in CSD. In (Desroches et al., 2019), the reversal potentials of the conductances of the GABAergic neuron did not depend on the ion concentrations, and only some of the transmembrane currents of the pyramidal neuron had an effect on ion concentrations and reversal potentials. Here, we took into account the effects of all the transmembrane currents on the intracellular ion concentrations of the respective neuron and on the extracellular ion concentrations. We replaced the Wang-Buzsáki model of the interneuron used in (Desroches et al., 2019) with a more recent model that better models features of fast-spiking cortical interneurons (Golomb et al., 2007). Moreover, we modified the GABAergic neuron leak current implementing sodium (0.012 mS cm*^−^*^2^) and potassium (0.05 mS cm*^−^*^2^) leak conductances, so that the GABAergic neuron does not spike in the absence of external input and its resting potential is in the physiological range.

Compared to (Desroches et al., 2019), we set the ratio of the GABAergic neuron/pyramidal neuron volume to 2/3, and, to include the impact of excitatory synaptic currents on the dynamics of ion concentrations, we separated the sodium and potassium components, assuming an equal permeability of the glutamatergic receptors to both ions. We modeled external inputs (drives) to the neurons using constant glutamatergic currents (which did not depend on the glutamate released by the synapses of the modeled neurons), whose amplitude was set by varying their conductances: *gD,e* for the pyramidal neuron and *gD,i* for the GABAergic neuron. We included the activity of the Na/K ATPase in the dynamics of ion concentrations for both neurons, not only for the pyramidal one as in (Desroches et al., 2019). We replaced the expression describing its dependence on the intracellular sodium and extracellular potassium concentrations with a more realistic one developed by (Kager et al., 2000), which is based on experimental data, and we introduced a voltage dependence of the pump function as in (Bouret et al., 2014), to prevent the membrane potential from reaching excessively negative values when recovering from a depolarization block. We used the half activation concentration values from (Huguet et al., 2016) and a maximal pump rate of 30 *µ*A cm*^−^*^2^ at *−*70 mV. We also modified a number of additional minor aspects. First, we increased the maximal conductance of the sodium leak current of the pyramidal neuron to 0.015 mS cm*^−^*^2^, so that its resting membrane potential is not overly negative. We also set the maximal conductance of the calcium-activated potassium current of the pyramidal neuron to 1 mS cm*^−^*^2^, the maximal conductance of the inhibitory synaptic current to 0.1 mS cm*^−^*^2^ and the rate of extracellular potassium diffusion to 0.00004 ms*^−^*^1^. For consistency, in the equation of the intracellular calcium concentration of the pyramidal neuron, we used the same factor as for the other ions to convert current density to rate of change in intracellular concentration.

We implemented the effect of Hm1a and of FHM3 mutations by replacing part of fast inactivating sodium current of the GABAergic neuron (modeled with a Hodgkin & Huxley) formalism with a persistent sodium current, keeping the sum of their maximal conductances constant; 1 % of persistent current was considered as the control physiologic condition. The equation modeling the persistent sodium current was similar to the one for the fast inactivating one, with the difference that there was no inactivation (we removed the h variable) and that the voltage dependence of the activation was shifted to more negative potentials by 12 mV (Bean, 2007; Magistretti and Alonso, 1999).

### Statistical analysis

For experiments with mice, we used data pooled from at least 3 animals per condition (including negative controls) to ensure reproducibility of results. Mice were used after genotyping and littermates were negative controls; in mice expressing fluorescent proteins, systematic tests using a flashlight were performed before the experiment to confirm the genotyping results. Statistical tests were performed with Origin 8 (OriginLab) and R. Fisher’s exact test followed by a pairwise test adjusted with Bonferroni correction was performed for the analysis of contingency tables. Two-tailed nonparametric Mann–Whitney *U test*, Kruskal-Wallis test and Friedman ANOVA were performed because groups are not large enough to accurately verify normality and equality of variance. Dunn’s nonparametric comparison and Bonferroni correction (comparisons with the control group) were used for post hoc tests when appropriate. The one sample Wilcoxon signed-rank nonparametric one-sided test was used to evaluate the effect of Hm1a on Na^+^ currents in cell lines (fold increase of I_NaP_ larger than 0) and the paired Wilcoxon Signed Rank Test was used for effect of Hm1a on firing in brain slices. Differences were considered significant at *p*<0.05.

## Results

### Acute gain of function of Na_V_1.1 channels induces CSD selectively in the neocortex

To disclose the role of Na_V_1.1 channels’ gain of function in the generation of CSD, we used the spider toxin Hm1a that has been reported as a specific Na_V_1.1 enhancer (Osteen et al., 2016). With whole-cell patch-clamp recordings of Na^+^ currents in cell lines, we first confirmed that also in our conditions Hm1a at low concentration (10 nM) selectively targets Na_V_1.1 over the two other Na_V_ isoforms expressed in the adult cortex, Na_V_1.2 and Na_V_1.6 (Supplementary Fig.S1). Importantly, Hm1a induced a 12-fold increase of persistent current, effect that is comparable to that previously observed with FHM3 mutations (Barbieri et al., 2019; Bertelli et al., 2019; Cestele et al., 2008; Cestele et al., 2013b; Dhifallah et al., 2018), making it a good pharmacological tool for modeling the effect of these mutations.

We tested the effect of the toxin in whole brain slices that included different structures, performing extracellular local field potential (LFP) recordings and intrinsic optical signal (IOS) imaging in an extended area (Fig.1). Bath application of 10 nM Hm1a lead to spontaneous CSD ignition, which was observed both as propagating wave in IOS images (Fig.1b_1_-b_2_) and as DC shift in LFP recordings (Fig.1c). CSD was elicited in 22.5% of the slices within 10min of Hm1a application (our time limit for determining successful induction), whereas we have never observed CSD in control conditions (Fig.1d). Interestingly, CSD was elicited only in the neocortex and never in the other structures monitored (hippocampus, dorsal striatum, globus pallidus and thalamus) (Fig. 1b_1_-b_2_ & d). To confirm that this was a neocortex-specific effect of Hm1a, we evaluated if in our conditions other brain areas were able to generate spreading depression by applying short puffs of 130mM KCl, a classic method of CSD induction (Pietrobon and Moskowitz, 2014; Zerimech et al., 2020). The success rate for CSD induction was 100% in all the structures tested: the neocortex (Supplementary Fig.S2a-c), the striatum (Supplementary Fig.S2d_1_) and the hippocampus (Supplementary Fig.S2d_2_). Neither Hm1a-induced nor KCl-induced CSD propagated outside the structure in which they were induced.

**Figure 1.**
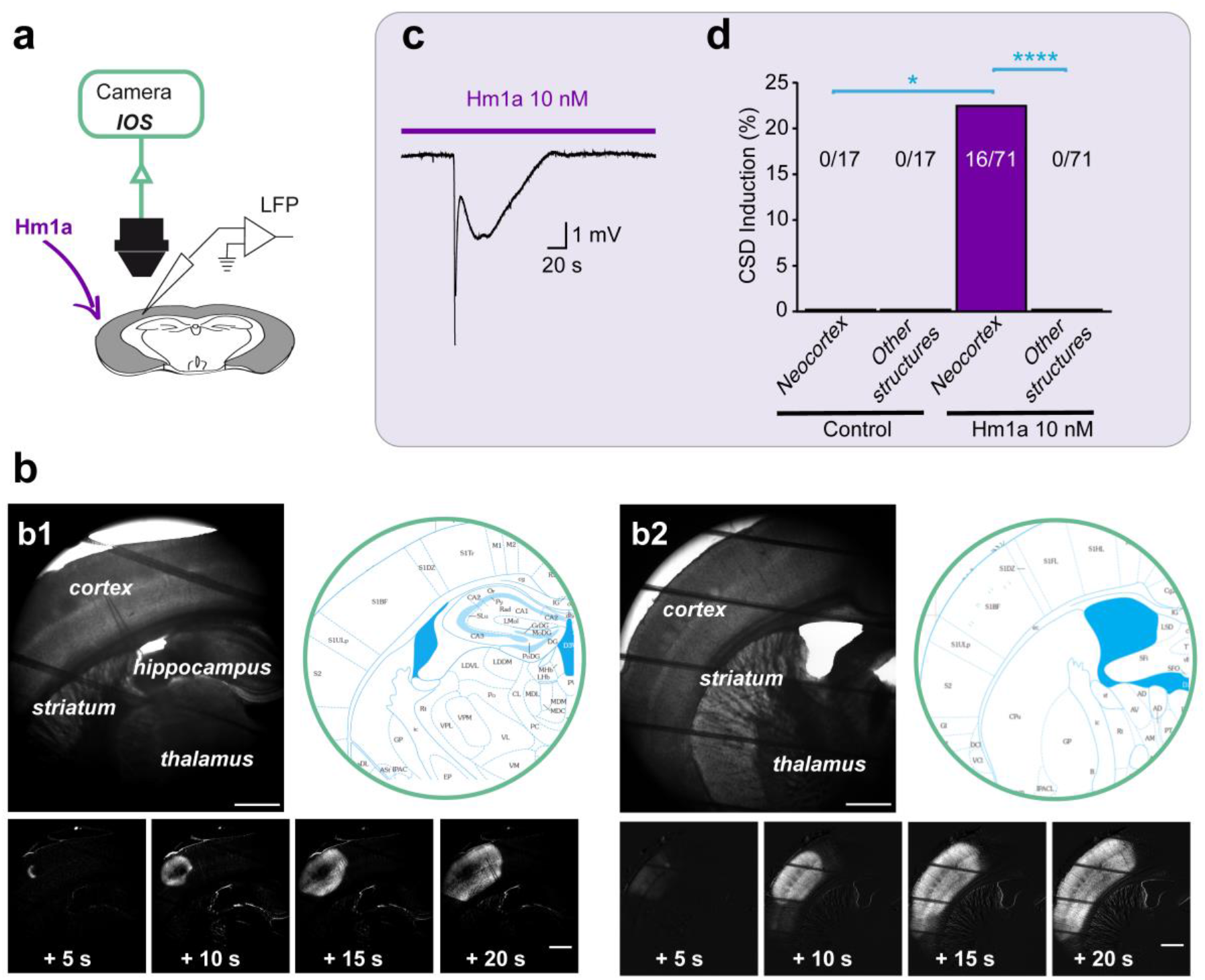
CSD can be triggered specifically in the neocortex by application of the selective Na_V_1.1 enhancer Hm1a. **a.** Experimental setting for studying CSD induction by Hm1a: brain slices were perfused with 10nM Hm1a, a concentration at which Hm1a is selective for Na_V_1.1 (see Suppl.Fig.1), and CSD induction was monitored with extracellular local field potential (LFP) recordings and intrinsic optical signal (IOS) imaging (obtained in extended brain regions with a 4X objective and a 0.35X camera adapter). **b_1_**. Upper left panel, transmitted light image of a representative coronal slice including the neocortex, the hippocampus, the dorsal striatum, the globus pallidus and the thalamus, as illustrated by the upper right panel obtained from Paxinos and Franklin’s Mouse Brain in Stereotaxic Coordinates 3^rd^ edition (Bregma 1.46, Interaural 2.34mm), in which the imaged area is indicated by the circle. The four bottom panels correspond to time series of image processing of IOS acquisitions (one image every 5 s, the first one 5 s after CSD initiation; see Methods), which show that CSD was triggered only in the neocortex. Scale bars: 500 μm. **b_2_**. Upper left panel, transmitted light image of another representative coronal slice including the neocortex, the dorsal striatum, the globus pallidus and the thalamus, as illustrated by the upper right panel obtained from Paxinos and Franklin’s Mouse Brain in Stereotaxic Coordinates 3^rd^ edition (Bregma -0.82, Interaural 2.98mm), in which the imaged area is indicated by the circle. The four bottom panels are time series of processed IOS images, which show that CSD was triggered only in the neocortex. Scale bars: 500 μm. **c**. Representative LFP recording of a CSD observed in the neocortex during the perfusion with Hm1a. **d.** Overall results showing the lack of spontaneous CSD in control (0/17 slices), and success rate for neocortical induction with bath-application of Hm1a (16/71), which never triggered CSD in other structures (Fischer’s exact test with Bonferroni correction, neocortex control vs neocortex Hm1a, * p=0.04; neocortex Hm1a vs other structures Hm1a, **** p=10^−4^).

**Figure 2.**
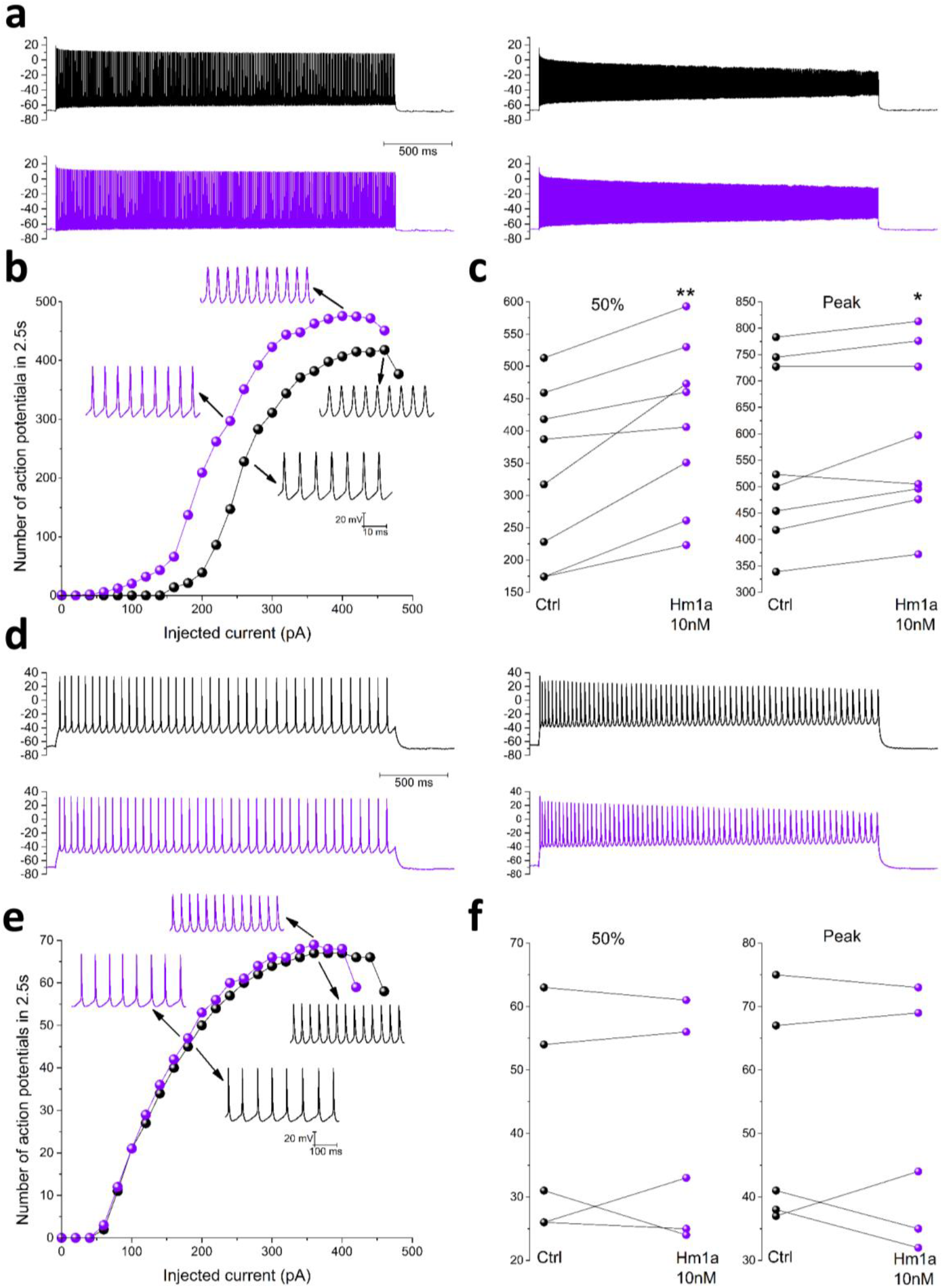
The selective Na_V_1.1 enhancer Hm1a increases the excitability of fast spiking GABAergic neurons but not of pyramidal neurons in neocortical brain slices. Recordings from neurons in neocortical Layer II-III of GAD67-GFP knock-in mice (which selectively label GABAergic neurons). **a.** Left, representative traces at 50% of the input-output relationship, recorded from a fast spiking neuron before (black) and after 10min (violet) perfusion with 10nM Hm1a; right, representative traces recorded from the same interneuron at the peak of the input-output relationship. **b.** Representative plot of the effect of Hm1a on the input-output relationship recorded from the same fast spiking interneuron; the arrows indicate magnified traces taken from a 50ms time window at the middle of the traces displayed in a, showing the increase in firing frequency induced by Hm1a at 50% and at the peak of the input-output relationship. **c.** Plot showing the change in firing frequency for each recorded neuron before and during Hm1a perfusion at 50% (28.4±7.0% mean increase, ** p=0.008) and at the peak of input-output relationship (7.1±2.6% mean increase, * p=0.03) (n=8). **d.** Left, representative traces at 50% of the input-output relationship, recorded from a pyramidal neuron before (black) and during (violet) perfusion with Hm1a; right, representative traces recorded from the same pyramidal neuron at the peak of the input-output relationship, before (black) and after 10min (violet) perfusion with Hm1a. **e.** Representative plot of the input-output relationship recorded from the same pyramidal neuron; the arrows indicate magnified traces taken from a 50ms time window in the middle of the traces displayed in d, showing that Hm1a has no effect. **f.** Plot showing the firing frequency for each recorded neuron before and during Hm1a perfusion at 50% (p=1) and at the peak of the input-output relationship (p=0.9). Wilcoxon Signed Ranks test for all the comparisons.

Then, we performed current-clamp patch-clamp recordings in brain slices from GAD67-GFP knock-in mice (which selectively label GABAergic neurons), to confirm that the Hm1a-induced gain of function of Na_V_1.1 can increase excitability preponderantly of GABAergic neurons in the neocortex. The resting membrane potential was maintained at around −70mV and action potential discharges were elicited with injections of 2.5s-long depolarizing current steps of increasing amplitude, comparing the properties of input output relationships before and after 10min perfusion with 10nM Hm1a. In order to minimize the variability of firing patterns of GABAergic neurons, we selected for the analysis fast-spiking neurons (Fig.2a-c). All the recorded pyramidal neurons showed regular spiking discharges (Fig.2d-f). The application of Hm1a induced in GABAergic neurons a leftward shift of the input-output relationship (28% mean increase of firing frequency at 50% of the input-output) and an increase of the maximal firing frequency (7% mean increase), whereas the firing properties of pyramidal glutamatergic neurons were not modified.

Thus, Hm1a induces in the neocortex hyperexcitability of GABAergic neurons but not of pyramidal neurons, and specifically triggers CSD in the neocortex, although in our experimental conditions spreading depolarizations can be induced by puffs of KCl and can be monitored by IOS imaging and LFP recordings in all the structures tested (neocortex, hippocampus and striatum).

### Optogenetic over-activation of GABAergic neurons can initiate CSD selectively in the neocortex

To directly investigate the role of GABAergic neurons, we used hemizygous VGAT.cre-ChR2(H134R)tdTomato.lox (VGAT-ChR2) mice, in which channelrhodopsin-H134R is selectively expressed in GABAergic neurons in different brain regions (Supplementary Fig.S3) (Chao et al., 2010). We activated GABAergic neurons in coronal brain slices illuminating with blue light an entire cerebral hemisphere, and we monitored CSD generation by both LFP recordings and IOS imaging of several brain structures (Fig.3a-b). Notably, similar to the experiments performed with Hm1a, the optogenetic activation of GABAergic interneurons induced CSD only in the neocortex: we never observed it in the hippocampus, striatum or thalamus. Mean latency to induction was 26.2 ± 1.8 s (n = 104) and propagation speed was 3.2 ± 0.1 mm/min (n = 104) (Fig.1c). Macroscopic features of CSD induced by optogenetic stimulation were similar to those of CSD triggered by a focal puff of 130 mM KCl in the neocortex (Supplementary Fig.S2b-c) or by perfusion with Hm1a (Fig 1b_1_-b_2_). We performed a subset of experiments for determining the rate of success of optogenetic CSD induction (Fig.3d; see methods for detailed procedure), observing that it was induced in 85 % of VGAT-ChR2 slices (always only in the neocortex) and never with control littermates (i.e. slices from VGAT-Cre, ChR2.lox or WT mice). Notably, CSD was readily induced by focal puff of 130 mM KCl in all the slices, both from VGAT-ChR2 mice and control littermates (Supplementary Fig.S2c), confirming that all the slices could generate CSD (not only those from VGAT-ChR2 mice). Moreover, as already highlighted before, CSD was readily induced by focal puff of 130 mM KCl also in the hippocampus and the striatum (Supplementary Fig.S2d). Thus, CSD initiation by over-activation of GABAergic neurons is a neocortex-specific mechanism, similarly to CSD initiation by Hm1a.

**Figure 3.**
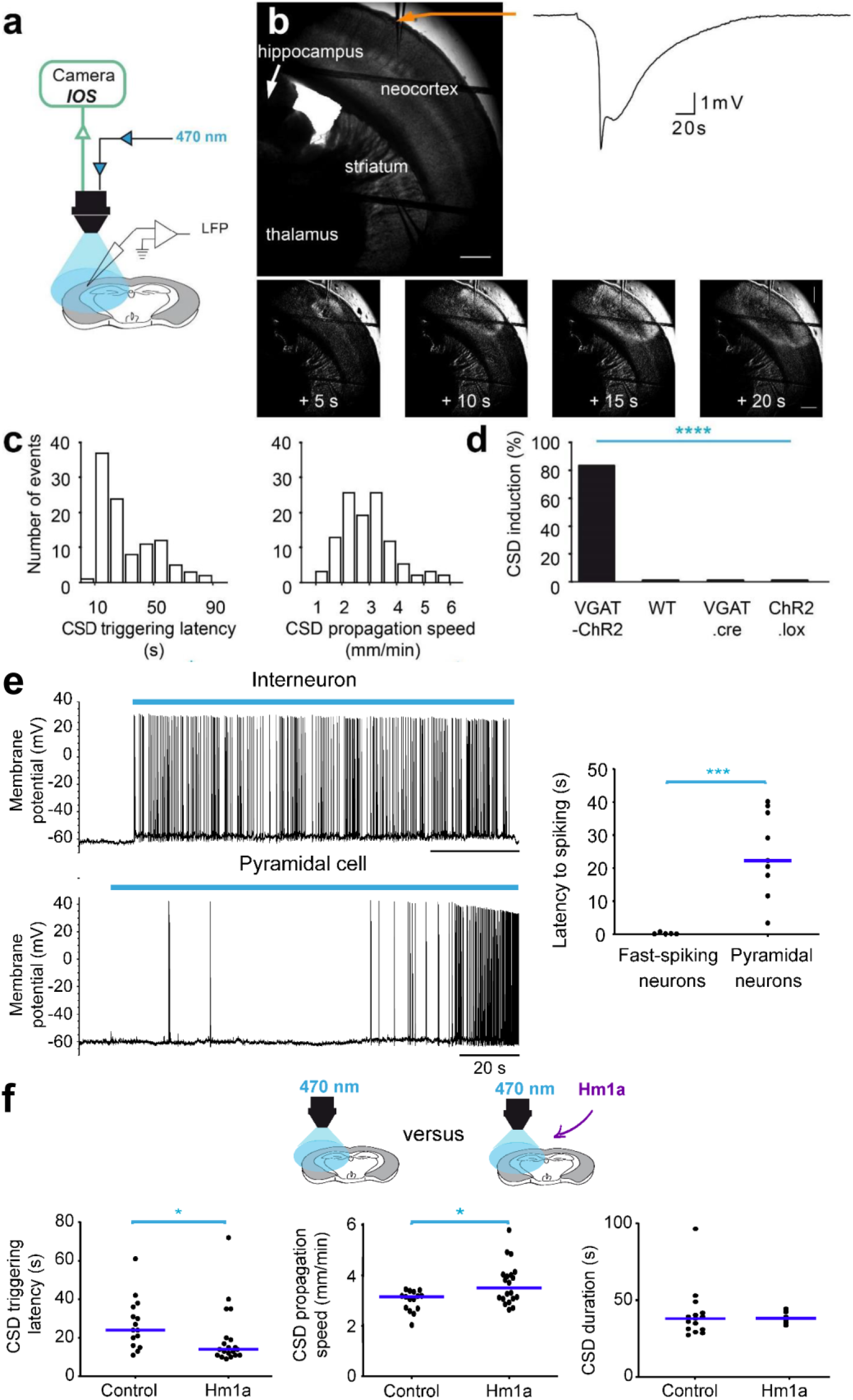
CSD is triggered specifically in the neocortex by optogenetic–induced hyperactivity of GABAergic neurons. **a.** Optogenetic stimulations were performed applying to coronal brain slices 470nm blue light, which was delivered with the 4x objective, illuminating a complete hemisphere (the area of illumination was larger than the area of image acquisition); CSD was measured with both extracellular LFP recordings and IOS imaging. **b**. Representative CSD that was induced only in the neocortex by the 470nm illumination in slices from VGAT-ChR2 mice (which express ChR2 selectively in GABAergic neurons), revealed by both the negative DC shift in the LFP and the IOS propagating wave. The four bottom panels are time series corresponding to image processed IOS acquisitions (one image every 5s, the first one 5s after CSD initiation; see methods). Scale bars: 500 μm. **c.** Left panel, distribution of latencies of CSD initiation upon 470nm illumination in VGAT-ChR2 slices (median=19 s, n=103 slices); right panel, distribution of propagation speed of optogenetic-induced CSD in VGAT-ChR2 slices (median=3.18 mm/min, n=103 slices). **d.** Success rate of optogenetic CSD obtained in a different series of experiments comparing slices from VGAT-ChR2 (11/13 slices), WT (0/14), VGAT.Cre (0/10) and ChR2.lox (0/10) mice (Fischer’s exact test, **** p=7 10^−5^). **e.** Left panels, representative whole-cell patch-clamp recordings of GABAergic and pyramidal neurons in layer II-III upon optogenetic illumination: a fast spiking GABAergic neuron responded to the 470nm illumination (blue bar) with short latency (upper panel), whereas a pyramidal neuron responded to the 470nm illumination (blue bar) with long latency (lower panel), scale bars=20 s; right panel, overall latencies to spiking during 470nm illumination for fast spiking neurons (median=0.30 s; n=5) and pyramidal neurons (median=22.26 s; n=12) (Mann-Whitney test, p=0.001); these recordings were not performed at the site of initiation, which with the large field of illumination used in these experiments was variable within the neocortex and not identifiable *a priori*. **f.** Further series of experiments in which features of optogenetic CSD induction were compared in VGAT-ChR2 slices perfused with Hm1a (but in which the toxin did not induce CSD within 10min) and control VGAT-ChR2 slices, waiting 10min before to illuminate (these experiments have been included in Fig.1d); left panel, latencies of optogenetic CSD measured in control VGAT-ChR2 slices (median=24 s, mean±SEM=27.3±3.4 s; n=15 slices) and VGAT-ChR2 slices perfused with Hm1a (median=14 s, mean±SEM=19.8±3.4 s; n=20 slices) (Mann-Whitney test, p=0.014); middle panel, propagation speed of optogenetic CSD in the same slices (control VGAT-ChR2, median=3.14 mm/min, mean±SEM=3.0±0.1 mm/min, n=15 slices; VGAT-ChR2 slices perfused with Hm1a without CSD, median=3.50 mm/min, mean±SEM=3.6±0.2 mm/min, n=20 slices) (Mann-Whitney test p=0.025); right panel, duration of optogenetic-induced CSD measured at half width of the LFP DC shift (control VGAT-ChR2 slices, median=38.1 s, mean±SEM=41.1±4.4 s, n=15 slices, VGAT-ChR2 slices perfused with Hm1a but without CSD, median=38.3 s, mean±SEM=38.7±1.7 s, n=6 slices) (Mann-Whitney test, p=0.7).

We then evaluated the response of pyramidal and GABAergic neurons in Layer II-III of the neocortex to the optogenetic stimulation before the induction of CSD, performing whole-cell patch-clamp recordings (Fig.3e). We observed that GABAergic neurons directly responded with short latency to the illumination, which triggered high frequency firing, whereas pyramidal neurons did not respond directly to the illumination and begun to spike after few tens of seconds of illumination.

In a further series of experiments, we evaluated the effect of Hm1a on optogenetic-induced CSD, hypothesizing a synergic role. In fact, optogenetic CSD was facilitated by Hm1a (Fig.3f): in brain slices in which Hm1a application did not induce CSD, subsequent optogenetic stimulation induced a 28% reduction of triggering latency and a 20% increase of propagation speed, compared to control slices; the duration of CSD evaluated measuring the LFP half-width was not modified by Hm1a. CSD was triggered only in the neocortex also in this series of experiments. Therefore, these results confirm the key role of Na_V_1.1 channels’ gain of function and over-activation of GABAergic neurons in the mechanism of CSD initiation that we have disclosed.

### Computational model of CSD initiation by overactivation of Na_V_1.1 channels and GABAergic neurons

We have recently shown that, in simulations obtained with a conductance-based model of a GABAergic neuron connected to a pyramidal neuron, an overactivation of the GABAergic neuron can lead to depolarizing block of the pyramidal neuron, which we considered the initiation of CSD (Desroches et al., 2019). In that model, the overactivation of the GABAergic neuron was obtained increasing its external depolarizing input (modeling a pathological state, as well as a condition similar to our optogenetic stimulation). Several putative GABAergic activation-related mechanisms were tested, and we identified the frequency of interneuron firing and the related increase of [K^+^]_out_ as the key element for inducing depolarizing block of the pyramidal neuron (Desroches et al., 2019).

Here, we have refined the model, in particular improving the spiking features of the GABAergic neuron and including complete dynamics of ion concentrations for both neurons (Fig.4a; see methods), so that the modifications of ion concentrations can modulate the activity of both neurons. The direct overactivation of the GABAergic neuron, with conditions similar to those that we used in (Desroches et al., 2019), generated simulations that were similar to those obtained in (Desroches et al., 2019) (not shown). Then, we tested the effect of an increase of the persistent Na^+^ current of the GABAergic neuron, which mimics the common effect of most FHM3 mutations (Barbieri et al., 2019; Bertelli et al., 2019; Cestele et al., 2013a; Cestele et al., 2008; Cestele et al., 2013b; Dhifallah et al., 2018; Fan et al., 2016; Mantegazza and Broccoli, 2019; Mantegazza and Cestele, 2017), as well as that of Hm1a. We modeled the control physiologic condition implementing a persistent conductance equal to 1% of the maximal sodium conductance, and we increased it up to 6% to simulate a pathological condition or the presence of Hm1a. In the physiological condition (Fig.4b), an external depolarizing input to the GABAergic neuron (*gD,i*) of 0.15 mS cm^−2^, without application of external input to the pyramidal neuron (*gD,e*), was able to induce high frequency firing of the GABAergic neuron. As in (Desroches et al., 2019), long lasting high frequency firing of the GABAergic neuron overcame its inhibitory effect, because it increased [K^+^]_out_. In the simulation of Fig.4b, [K^+^]_out_ reached a maximum of 10.6 mM, depolarizing the pyramidal neuron and triggering its spiking, which was transient and ended after few seconds, when [K^+^]_out_ relaxed to lower levels. In this condition, there was no depolarizing block (CSD initiation). When the same simulation was run with persistent sodium conductance of the GABAergic neuron increased to 6% (Fig.4c), in the initial phase the GABAergic neuron discharged at higher frequency leading to a larger increase of [K^+^]_out_ (up to 13.9 mM). Importantly when the pyramidal neuron was engaged in firing, both neurons underwent depolarizing block, with [K^+^]_out_ rising to >60 mM, indicating CSD initiation. In fact, the increase of persistent sodium current of the GABAergic neuron facilitated CSD, leading to a decrease of the minimal *gD,i* that can induce CSD initiation (Fig.4d), Moreover, similar to the experimental data of Fig.3f, the increase of persistent current lead to a reduction of the CSD initiation latency (Fig.4e); CSD was induced in this simulation with *gD,i*= 0.395 mS cm^−2^, which was the minimal input able to induce CSD with 1% persistent current, see Fig.4d).

**Figure 4.**
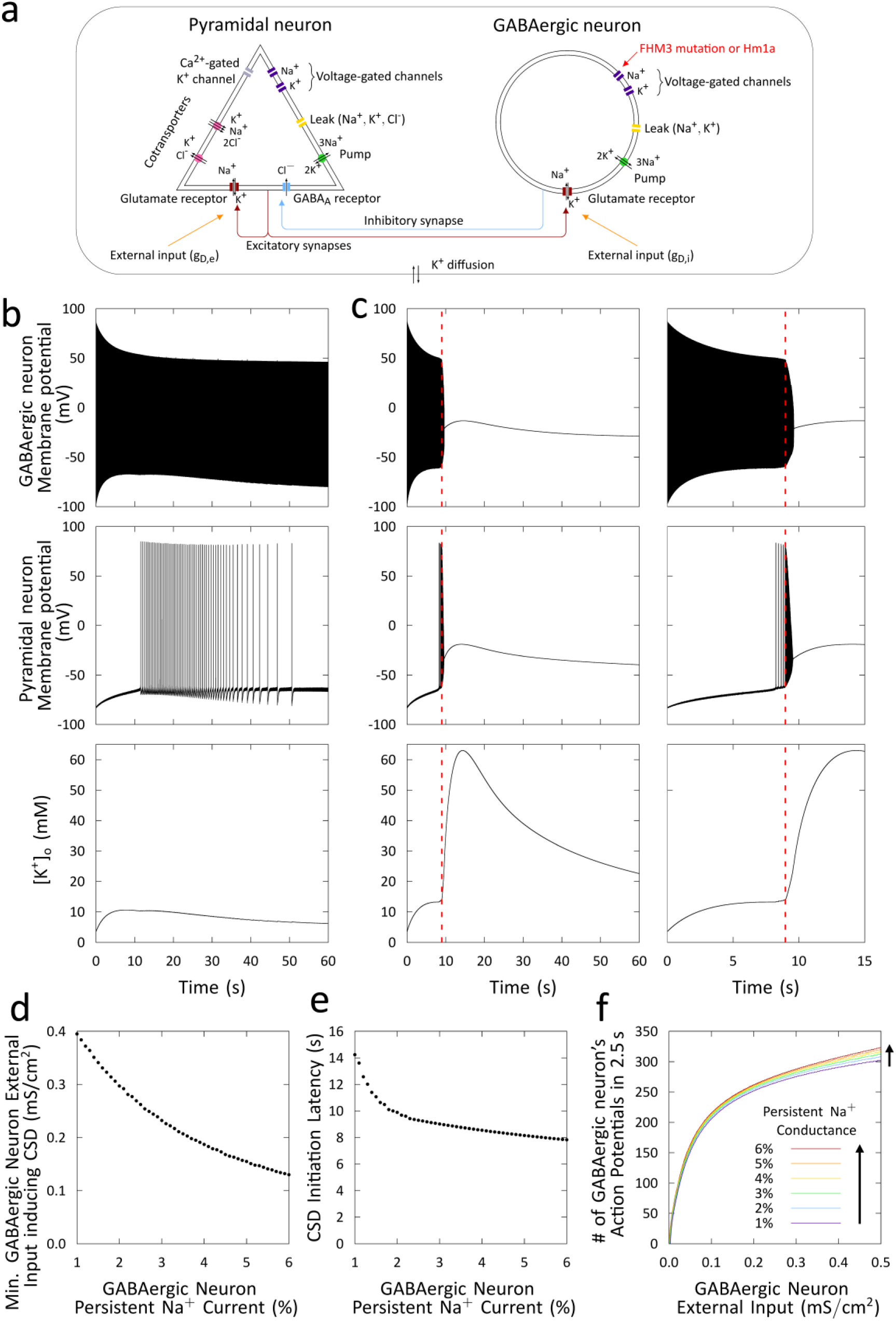
The increase of GABAergic neuron’s persistent current and the consequent hyperexcitability facilitates the initiation of CSD in a computational model. **a.** Diagram illustrating the conductance-based computational model that we developed extending and improving the model of a coupled GABAergic-pyramidal neuron pair that we used in (Desroches et al., 2019). Here, a number of modifications have been implemented, including a more realistic GABAergic neuron and a complete modeling of the dynamics of ion concentrations for both neurons (see methods). We modeled the effect of migraine mutations and of Hm1a by increasing the persistent sodium current of the GABAergic neuron. *gD,e* and *gD,i* are glutamatergic conductances that model the baseline excitatory inputs (“excitatory drive”) of the pyramidal and of the GABAergic neuron, respectively. **b**. Simulation, with GABAergic neuron’s physiologic persistent sodium current (1%), of the effect of a constant external depolarizing input to the GABAergic neuron (*gD,i* = 0.15 mS cm^−2^) without input to the pyramidal neuron (*gD,e* = 0 mS cm^−2^): membrane potential of the GABAergic neurons (upper panel), membrane potential of the pyramidal neuron (middle panel) and extracellular potassium concentration (lower panel). **c**. Same simulation with increased GABAergic neuron’s persistent sodium current (6%, mimicking the effect of FHM3 mutations and of Hm1a); the right panels display with an enlarged time scale the first phase of the simulation shown in the panels on the left. The vertical red line is the beginning of the large [K^+^]o increase that leads to depolarizing block. **d**. Effect of an increase of the GABAergic neuron’s persistent sodium current on the lowest external input to the GABAergic neuron (*gD,i*) sufficient to induce depolarizing block. **e**. Effect of an increase of the GABAergic neuron’s persistent sodium current on the depolarizing block latency with *gD,i*=0.395 mS cm^−2^ (the lowest external input to the GABAergic neuron able to generate CSD in control conditions: 1% persistent sodium conductance). **f**. Effect of the level of persistent current on the firing frequency of the GABAergic neuron, in a simulation in which the pyramidal neuron has been removed, reflecting the direct effect of the persistent current on the firing properties of the GABAergic neuron.

Finally, we evaluated, in simulations in which the pyramidal neuron was removed from the model, the effect of an increase of persistent current on the “intrinsic” firing frequency of the GABAergic neuron, observing that it was increased at all the levels of external input tested (Fig.4f), showing that the persistent current can increase the firing frequency of the GABAergic neuron independently from the interactions with the pyramidal neuron (e.g. synaptic input, modifications of ion concentrations).

Overall, the model shows that an increase of persistent sodium current, mimicking FHM3 mutations or Hm1a, induces hyperexcitability of the GABAergic neuron, leading to a facilitation of CSD, which is ignited with lower values of external input and shorter latency, consistent with the experimental data. Interestingly, a prediction of the model obtained in the simulations presented here and in those of (Desroches et al., 2019), is that the key factor for CSD initiation induced by overactivation of GABAergic neurons is an increase of [K^+^]_out_, initially directly generated by the spiking of the GABAergic neuron. Thus, we investigated in our experimental system the detailed mechanisms of CSD initiation.

### Detailed mechanism of CSD initiation

We initially evaluated the effect of Na_V_1.1 loss of function on CSD initiation, to compare it with the gain of function tested above. We crossed VGAT-ChR2 mice with knock-out *Scn1a*^+/−^ mice, which model the epileptic encephalopathy Dravet syndrome and in which one allele of the *Scn1a* gene is not functional, causing Na_V_1.1 haploinsufficiency and hypo-excitability of GABAergic neurons (Mantegazza and Broccoli, 2019; Yu et al., 2006). This selective effect on GABAergic neurons has been observed in numerous studies, also at the age that we have used for our studies, although there could be remodeling at later developmental stages (Favero et al., 2018). In brain slices from VGAT-ChR2/ *Scn1a*^+/−^ mice, the success rate of optogenetic CSD induction was reduced 2.4-fold (Fig.5a and Supplementary Table), showing that Na_V_1.1 loss of function and reduced excitability of GABAergic neurons can inhibit CSD initiation by optogenetic activation of GABAergic neurons. Contrarily than the inhibition of initiation, there was a trend towards an increase of the propagation speed (Supplementary Table), consistent with different mechanisms of initiation and propagation.

**Figure 5.**
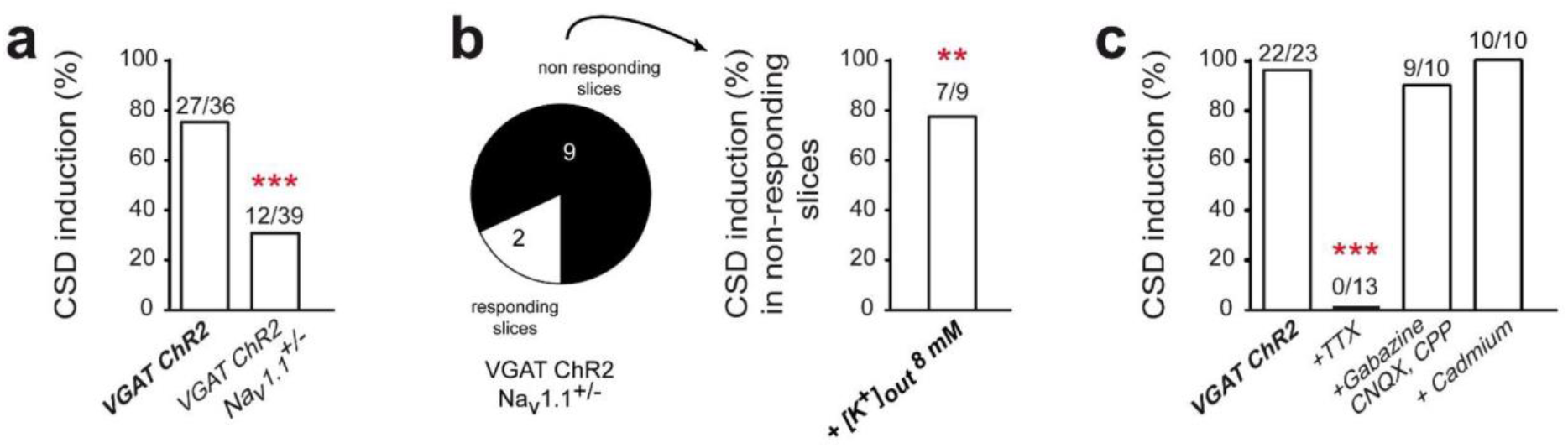
Effect of the reduction of GABAergic neurons’ excitability, block of neuronal excitability or block of synaptic transmission on optogenetic CSD induction. **a**. Reduction of the success rate of optogenetic CSD induction in slices from VGAT-ChR2-*Scn1a*^+/−^ mice compared to VGAT-ChR2 littermates (*** p=0.0002, Fischer’s exact test). **b**. A different series of experiments in which optogenetic CSD was induced after the increase of [K^+^]_out_ to 8mM in VGAT-ChR2-*Scn1a*^+/−^ slices in which optogenetic CSD was previously not induced with standard [K^+^]_out_ (*** p=0.002 Fischer’s exact test). **c**. Success rate of optogenetic CSD in VGAT-ChR2 slices perfused with a Na^+^ channel blocker (TTX-1 μM), with GABA-A and NMDA-AMPA-Kainate receptor antagonists (Gabazine-15μM, CPP-10μM, CNQX-20μM), or with a Ca^2+^ channel blocker (Cd^2+^ 100μM) to fully block synaptic release (Fisher’s exact test, **** p=1.4e-11; Bonferroni-corrected post-test, *** p<0.0001).

Then, we tested the importance of extracellular K^+^ accumulation as a key parameter for CSD initiation. We initially used VGAT-ChR2/*Scn1a*^+/−^ slices in which the optogenetic stimulation did not trigger CSD, perfusing them with mACSF in which [K^+^] was moderately increased (from 3.5 to 8 mM), and applying a second optogenetic stimulation. Notably, we found that the reduced success rate of CSD induction in VGAT-ChR2/*Scn1a* ^+/−^ slices was rescued with 8mM [K^+^]_out_ (Fig.5b).

Although our computational model points to extracellular K^+^ build-up directly induced by the spiking of GABAergic neurons, it has been shown that hyperactivity of GABAergic neurons can favor neuronal network excitation also by other mechanisms, including synaptic transmission-driven activation of neuron-glia networks (Kaila et al., 2014; Perea et al., 2016; Shiri et al., 2016). Thus, we performed pharmacological experiments to disclose the detailed mechanism linking hyperactivity of GABAergic neurons to CSD initiation. In particular, the K^+^-Cl^−^ cotransporter KCC2 can induce post-synaptic K^+^ efflux and is involved in excitatory actions of GABAergic transmission leading to hyperexcitability (Kaila et al., 2014). However, two different selective KCC2 inhibitors did not modify the success rate and dynamics (latency, propagation speed) of CSD induced by optogenetic stimulation, not even upon pre-treatment of slices with the GABA-A receptor agonist isoguvacine that we used to increase KCC2 baseline activity (Supplementary Table). Further, we tested blockers of neuronal excitability or synaptic transmission.

The Na^+^ channel/action potential blocker tetrodotoxin (TTX) completely suppressed CSD induction (Fig.5c), whereas the block of glutamate (Kainate-AMPA-NMDA) and/or GABA-A receptors (with CNQX-CPP and gabazine, respectively), or the complete block of synaptic transmission with the Ca^2+^ channel blocker Cd^2+^ (Fig.5c, Supplementary Table) did not modify the success rate of CSD induction. Notably, neurotransmission, in particular glutamatergic one, was instead important for sustaining CSD propagation, because in the presence of CNQX-CPP or Cd^2+^, the speed of CSD propagation was reduced and the propagation often aborted (Table). Altogether, these results demonstrate that GABAergic neurons’ hyperexcitability is sufficient for CSD initiation in the neocortex, that CSD is not directly triggered by ion flux through channelrhodopsin, and that synaptic transmission-driven mechanisms are not necessary, although they are implicated in propagation. Also, they suggest that the mechanism of initiation could involve spike-generated [K^+^]_out_ increase.

**Table.**
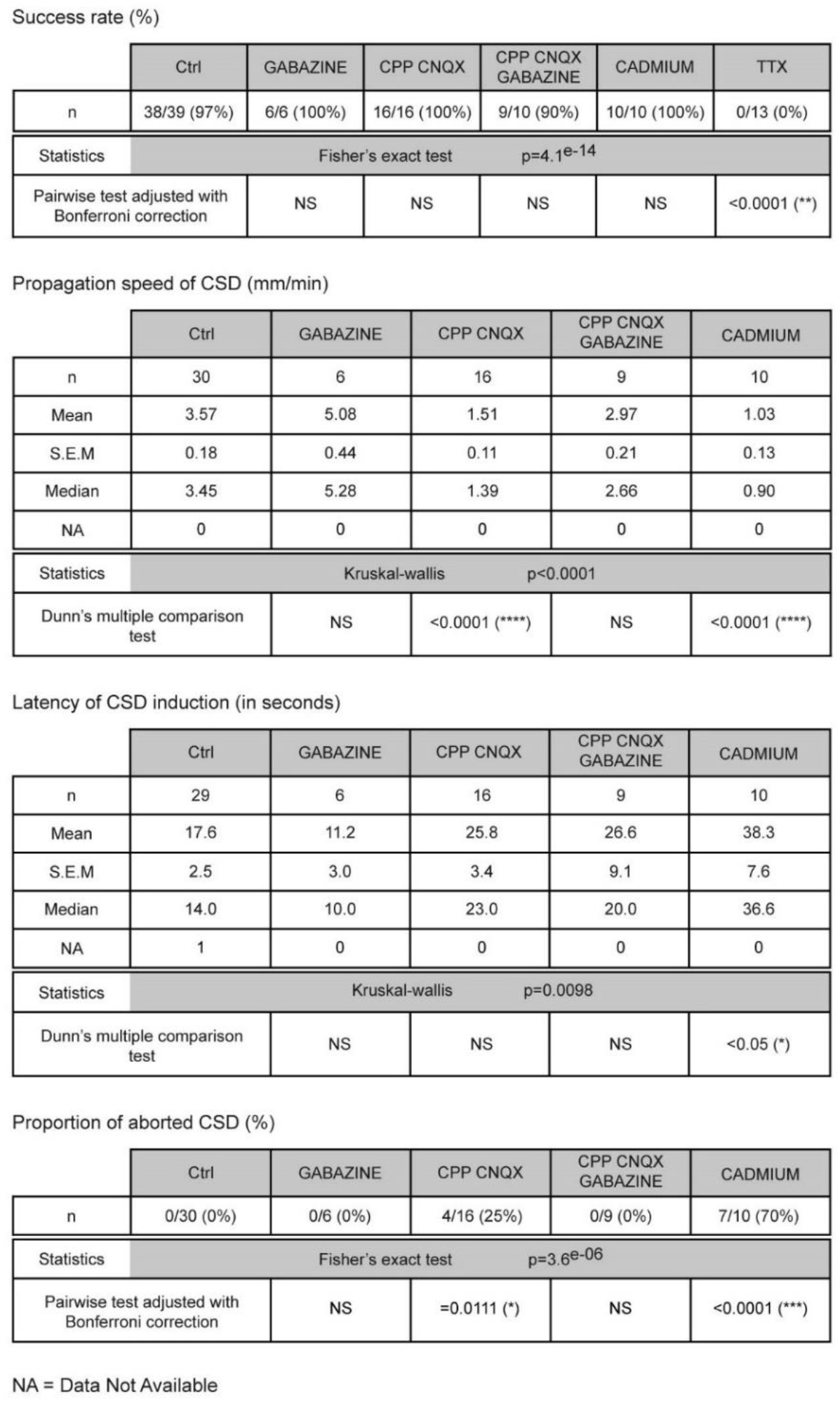
Effect of block of synaptic transmission or neuronal excitability on CSD induced by optogenetic stimulation. Success rate (%), propagation speed (mm/min) and initiation latency (seconds) of CSD induced 470 nm illumination in slices from VGAT-ChR2 mice treated with a GABA-A receptor antagonist (Gabazine-15 μM) and/or NMDA-AMPA-Kainate receptor antagonists (CPP-10 μM, CNQX-20 μM), with the calcium channel/synaptic release blocker cadmium (Cd^2+^-100 μM), or with the sodium channel/action potential blocker tetrodotoxin (TTX-1 μM). The number of CSD, induced by 470 nm illumination, that quickly stop after initiation (reported as aborted CSD) is also quantified. Note that some of these results have been already presented in Fig.4c. “n” refers to the number of slices, and “NA” to data not available (see methods).

To disclose whether spike-generated [K^+^]_out_ increase was directly involved in CSD initiation, we first evaluated the [K^+^]_out_ dynamics during optogenetic illumination, measuring [K^+^]_out_ at the site of CSD initiation with K^+^-sensitive electrode recordings, together with LFP/multi-unit activity (MUA) recordings. In order to perform recordings at the site of initiation, we used spatial optogenetic illumination to specifically control the site of CSD induction (Fig.6a-b). Our data shows that [K^+^]_out_ slowly and progressively increased during illumination, reaching ∼12 mM when CSD was ignited (identified as the end of the multi-unit activity) (Fig.6c-d). This suggests that, at CSD initiation, [K^+^]_out_ can progressively accumulate near neuronal membranes, until CSD ignition threshold is reached. We confirmed this hypothesis by inducing CSD with focal applications of 12 mM KCl in a neocortical area that was similar in size to that of spatial optogenetic illuminations (Fig.6e).

**Figure 6.**
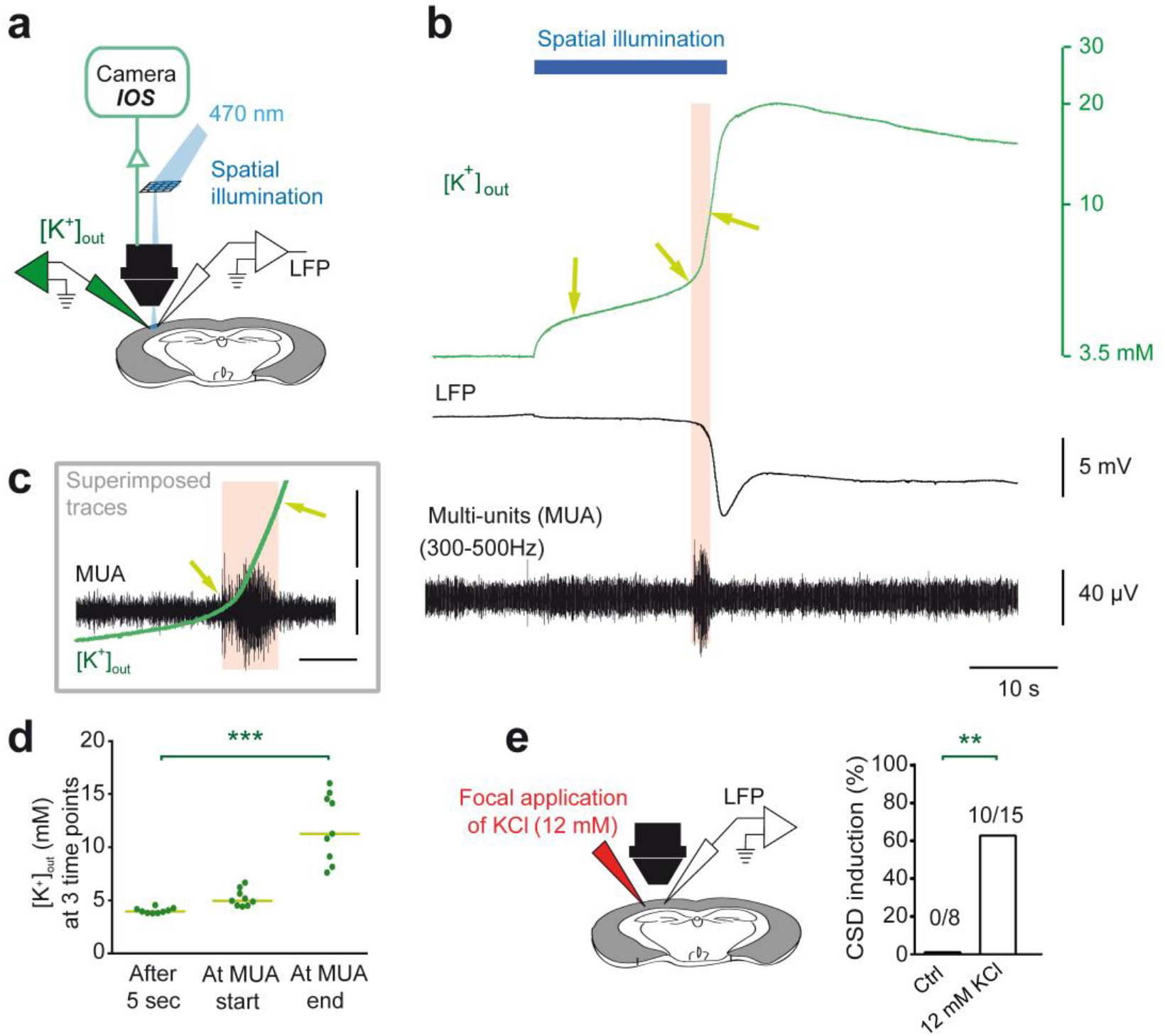
Spike-induced increase of [K^+^]_out_ is directly involved in CSD induction by spatial illumination. **a.** Experimental setting for spatial 470 nm illumination used to specifically control the area of CSD induction, allowing [K**^+^**]_out_, LFP and IOS recordings at the site of CSD initiation. **b**. [K**^+^**]_out_ dynamics before and during CSD, correlated to the LFP and multi-unit activities (MUA), which were paroxysmal at CSD initiation. Only the first component of the CSD is shown. **c.** Enlargement and superposition of [K**^+^**]_out_ and MUA traces shown in b. **d**. Quantifications of [K**^+^**]_out_ after the first 5sec of illumination, at the beginning of the paroxysmal MUA firing and at the end of the MUA firing (beginning of the depolarizing block) (arrows in b and c), n=9 slices. Bars represent medians. Friedman test (p<0.0001) and Dunn’s post-test (*** p<0.001). **e**. Success rate of CSD induced by long-lasting puff of 12mM KCl (dissolved in 125mM NaCl), which corresponds to the [K**^+^**]_out_ at the beginning of the depolarizing block, compared to a control 137mM NaCl solution (Ctrl; Fisher exact test, ** p=0.0027). Injection area: 0.75 +/− 0.11mm^2^ (n=10 slices with successful CSD inductions).

### Overactivation of GABAergic neurons can initiate CSD *in vivo*

Thus, to show that overactivation of GABAergic neurons can lead to CSD induction also *in vivo*, condition in which there is blood circulation, long range connections and neuromodulations, we illuminated with an optical fiber the somatosensory cortex of VGAT-ChR2 mice, monitoring CSD by LFP recordings (Fig.7); CSD was induced in 50 % of VGAT-ChR2 mice and never in control littermates. Therefore, this data confirms the results obtained in brain slices with optogenetic stimulations.

**Figure 7.**
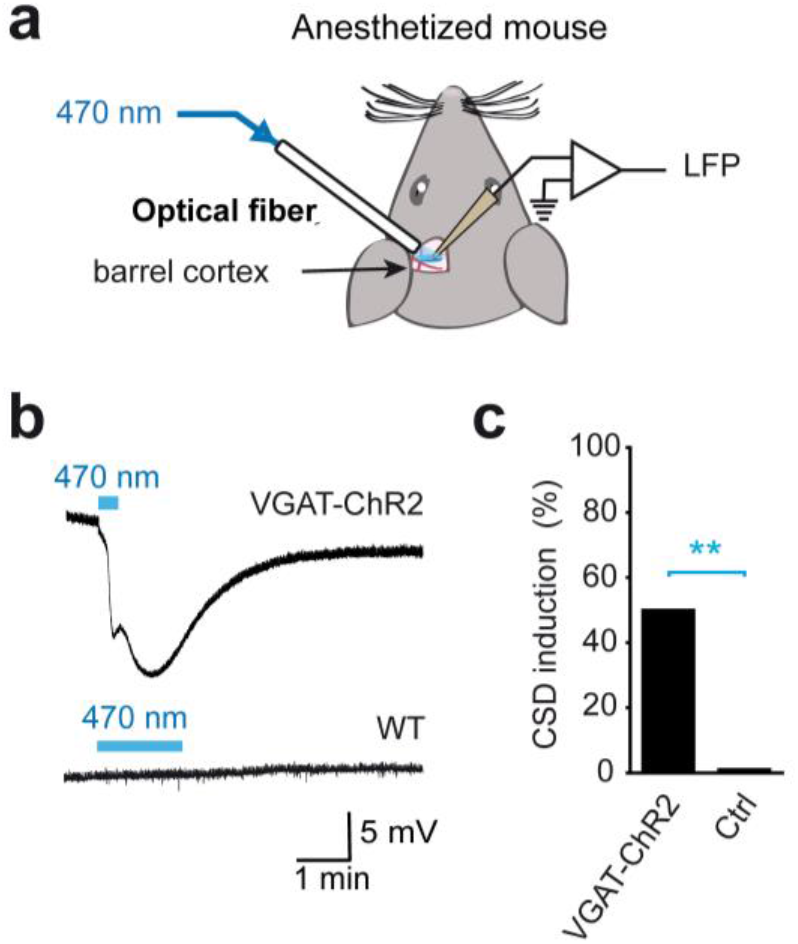
Optogenetic CSD induction *in vivo*. **a.** Experimental design: blue light optogenetic stimulation (100 Hz trains of 0.8 ms pulses) was applied to the barrel cortex of anesthetized mice with an optical fiber through a craniotomy; DC field potential recordings were performed with a glass pipette (Ag/AgCl electrode). **b.** Representative field potential traces of CSD in a VGAT-ChR2 mouse, whereas there was no response in a WT mouse. **c.** Proportion of optogenetic CSD induction in VGAT-ChR2 (5/10) and control mice (0/13, including WT, N = 5, VGAT.Cre, N = 4, and ChR2.lox N = 4. Fischer’s exact test; ** p=0.0075).

## Discussion

We have identified and characterized a novel mechanism of CSD initiation specific of the neocortex, showing in acute experimental models a causal relationship between initial hyperactivity of GABAergic neurons and CSD ignition, driven by the progressive increase of [K^+^]_out_ at the initiation site and in which synaptic transmission is not necessary (Fig.8). This mechanism was supported by simulations obtained with a computational model. CSD was induced in the neocortex both by optogenetic activation of GABAergic neurons and by activation of Na_V_1.1 with the specific toxin Hm1a (Osteen et al., 2016). This is consistent with the key role of Na_V_1.1 in GABAergic neurons’ excitability and with the effect of FHM3 mutations, which cause gain of function of Na_V_1.1 and can induce an increase of the persistent Na^+^ current similar to that observed with Hm1a (Barbieri et al., 2019; Bertelli et al., 2019; Cestele et al., 2013a; Cestele et al., 2008; Cestele et al., 2013b; Fan et al., 2016; Mantegazza and Broccoli, 2019; Mantegazza and Cestele, 2017). Notably, it has been recently reported that Hm1a rescued the hypoexcitability of hippocampal GABAergic neurons and the severity of the epileptic phenotype in epileptic *Scn1a*^+/−^ knock-out mice, but it did not modify firing properties of wild type hippocampal GABAergic neurons in brain slices (Richards et al., 2018). However, we found that application of Hm1a at a concentration at which it is specific for Na_V_1.1 can induce hyperexcitability of GABAergic neurons in neocortical slices from wild type mice, whereas the firing properties of glutamatergic pyramidal neurons were not significantly modified, consistent with the different role of Na_V_1.1 in the two neuron subtypes.

**Figure 8.**
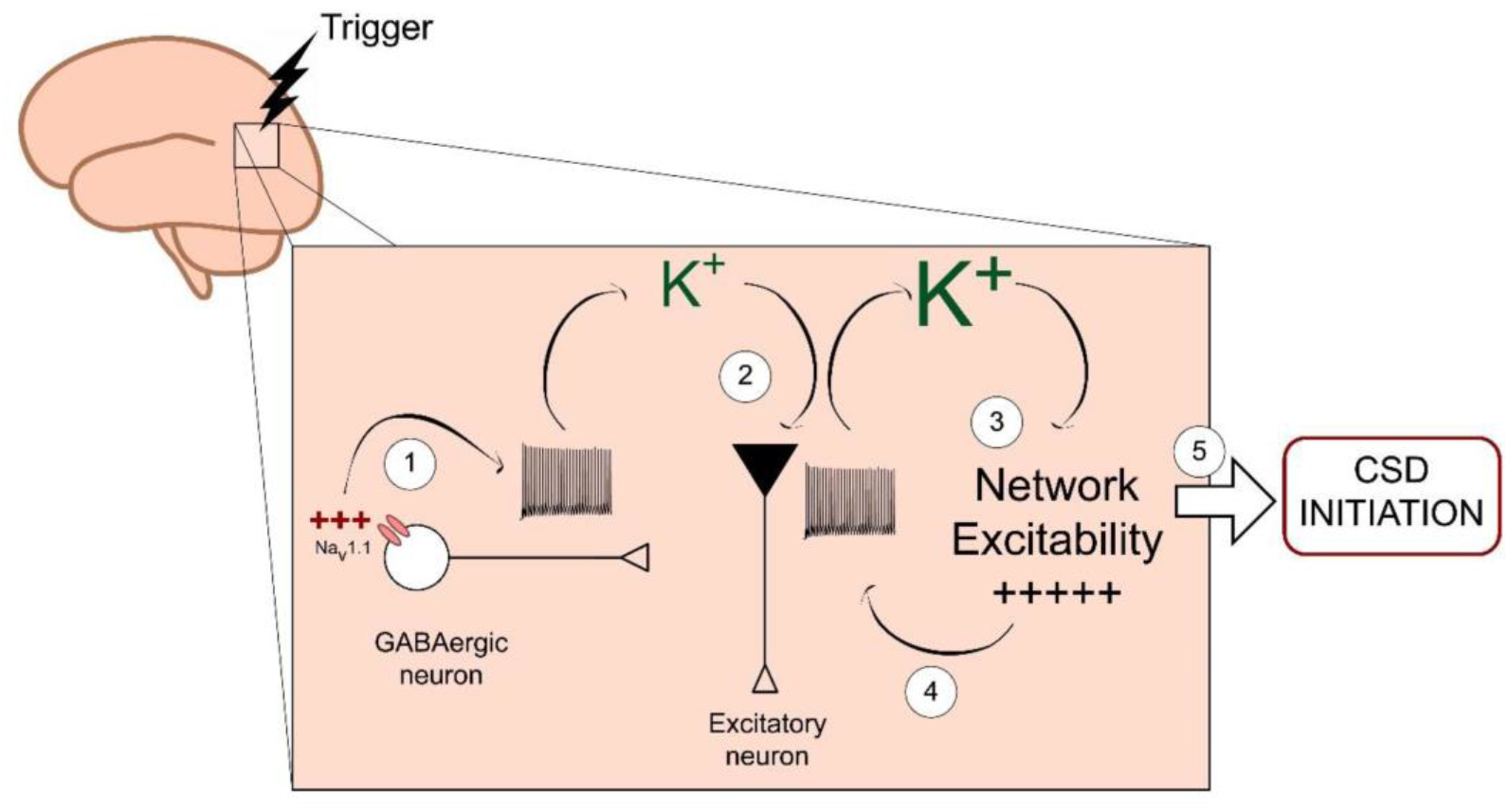
Diagram of the novel mechanism of CSD initiation. GABAergic neurons’ hyperactivity and intense firing (1) caused by pathologic dysfunctions (e.g. Na_V_1.1 gain of function) can lead to spiking-induced accumulation of extracellular K^+^ (2), which drives the network to hyperexcitability (3) and eventually induces depolarization block and CSD initiation, independently from synaptic transmission. This mechanism can be associated not only to gain of function Na_V_1.1 mutations, but possibly also to other dysfunctions that induce GABAergic neurons’ hyperactivity. As in all episodic disorders, homeostatic mechanisms can control dysfunctions in the period between the attacks, but triggering factors (e.g. hormonal/neuromodulatory changes or increase of incoming neuronal signals from the periphery) may focally affect neuronal excitability and activities of cortical networks, leading to long lasting GABAergic neurons’ hyperexcitability and CSD induction.

Additionally, we found that the loss of function of Na_V_1.1 in slices from VGAT-ChR2-*Scn1a*^+/−^ mice (Yu et al., 2006) inhibited the initiation of CSD by optogenetic stimulation, consistent with the involvement of Na_V_1.1 and of GABAergic neurons’ hyperexcitability in the mechanism of initiation. This could appear puzzling at first sight, because CSD is characterized by network hyperexcitability and cortical networks are hyperexcitable in mice carrying loss of function Na_V_1.1 mutations (Hedrich et al., 2014; Liautard et al., 2013). Moreover, it has been shown that Na_V_1.1 loss of function facilitates spreading depression in the brainstem, which can cause post-seizure sudden death (SUDEP) in *Scn1a*^+/−^ mice because of block of cardiorespiratory pacemaking (Aiba and Noebels, 2015). However, these apparent discrepancies could be consistent with different mechanisms of initiation and propagation. In fact, contrary to induction, we did not observe inhibition of CSD propagation in VGAT-ChR2-*Scn1a*^+/−^ slices, in which there is reduced excitability of GABAergic neurons, but we observed instead a trend towards facilitation. This could facilitate the generation/propagation of spreading depression in the brainstem upon induction of epileptic activity with post-ictal depression in the cortex, as observed In *Scn1a*^+/−^ mice by (Aiba and Noebels, 2015).

This further highlights the role of GABAergic neurons in the mechanism of CSD initiation that we have studied, and suggests that specific types of network hyperexcitability are required for different mechanisms of CSD initiation. Thus, GABAergic neurons’ hyperactivity can acutely induce CSD in the normal (non-pathologic) neocortex, and, in comparison with possible chronic models (i.e. genetic), this approach demonstrates that other pathological modifications (e.g. remodeling) are not necessary. This is a novel, previously undisclosed, role of GABAergic neurons in CSD.

We have also demonstrated that CSD initiation by direct GABAergic neurons’ hyperactivation is not dependent on synaptic transmission. The manipulations that inhibited CSD initiation were the generalized block of action potentials’ generation (applying TTX) or the reduction of GABAergic neurons’ excitability (in brain slices from VGAT-ChR2-*Scn1a*^+/−^ mice), consistent with the simulations that we obtained in (Desroches et al., 2019). Differently than initiation, CSD propagation was inhibited by blocking Ca^2+^ channel-dependent synaptic release or glutamate receptors, but not by blocking GABA-A receptors. These results show that the increase in GABA-A receptor activation caused by GABAergic neurons’ hyperexcitability is not involved in CSD, whereas glutamatergic transmission has an important role in propagation, but not in initiation when GABAergic neurons are directly overactivated. Overall, this data is consistent with different mechanisms of initiation and propagation. Optogenetic experiments with spatial illumination (Fig.6) allowed us to investigate the properties of the site of initiation. Numerous studies have performed pharmacological investigations of mechanisms of CSD triggered with classic methods, revealing a complex picture with results that often are specific for different methods and for initiation vs. propagation (Pietrobon and Moskowitz, 2014), consistent with different cellular/molecular mechanisms. Neuronal firing-induced K^+^ build-up is a key factor for CSD ignition by hyperactivation of GABAergic neurons, a novel mechanism of induction that is different in comparison with that implicated in models of FHM1 & 2, in which it has been proposed that excessive glutamate release/accumulation is the major pathological dysfunction (Capuani et al., 2016; Tottene et al., 2009; Vecchia et al., 2014). Notably, a recent optogenetic CSD model of glutamatergic neurons’ hyperactivation probably mimics mechanisms at play in FHM1 & 2, including the necessity of NMDA receptor activation for CSD initiation (Chung et al., 2018).

Consistent with a different mechanism in comparison with FHM3, FHM1 & 2 patients often show complex phenotypes that are more severe than those of FHM3 and include several neurologic/psychiatric co-morbidities, including seizures that herald or are concomitant with hemiplegic migraine attacks, as well as peri-ictal death (Ferrari et al., 2015; Vecchia and Pietrobon, 2012). In FHM3 there are no patients with these complex phenotypes, and in the few cases in which seizures have been reported, they are always independent from migraine attacks and present in different developmental windows (Mantegazza and Cestele, 2017). Moreover, migraine is not part of the phenotypes of epileptogenic Na_V_1.1 mutations, which cause loss of function of the channel and hypoexcitability of GABAergic neurons (Dravet et al., 2005; Zhang et al., 2017; Zuberi et al., 2011), consistent with different pathologic mechanisms of epileptogenic and migraine Na_V_1.1 mutations. Congruously, we have never observed ictal-like epileptiform activities in our experiments, although it has been shown that ictal-like activities generated by application of convulsants in brain slices could be enhanced/induced (Shiri et al., 2016; Yekhlef et al., 2015) or, depending on the brain region, inhibited (Ledri et al., 2014) by the optogenetic activation of GABAergic neurons. This reflects the requirement of a network that already generates epileptic activities for induction/modulation of these activities by the activation of GABAergic neurons. Nevertheless, it is interesting to note that, in some of those studies, different mechanisms of onset (some requiring activation of GABAergic neurons, others activation of glutamatergic neurons) have been identified for epileptiform activities that appear phenomenologically similar (Shiri et al., 2016). This could be the case, as we have shown, also for network activities that lead to CSD induction, which could be specific for different types of migraine, in particular for FHM3 compared to FHM1 & 2.

In our experiments, latencies to CSD induced by activation of GABAergic neurons were of few tens of seconds. In most of the classic experimental models, CSD is induced with strong stimuli (e.g. injections of KCl in the tens of mM to molar range, the latter well beyond pathophysiological limits) (Pietrobon and Moskowitz, 2014), which often result in shorter latencies most likely because they do not reproduce the complete dynamic process of CSD onset at the initiation site, mimicking conditions that are those of the generalized depolarization phase. There is no information about activities of single neurons leading to CSD induction in migraine patients. As in other episodic and paroxysmal disorders (Ptacek, 2015), pathologic dysfunctions of migraine are probably controlled by homeostatic mechanisms in the period between attacks, and different factors (e.g. hormonal/neuromodulatory changes or increase of incoming neuronal signals from the periphery) may affect neuronal excitability and activities of cortical networks, triggering CSD induction and migraine attacks. It can be hypothesized that, in the restricted volume of cortex in which CSD is initially ignited, these factors could weaken homeostatic controls and hyperexcitable GABAergic neurons could be hyperactivated for tens of seconds, similarly to our acute model. Interestingly, neurons can show early and long lasting increase of activity also in other episodic neurologic disorders, as observed with single unit recordings in epileptic foci of patients (Truccolo et al., 2011), in which increased neuronal activity begins minutes before the attack.

In conclusion, we have disclosed a novel mechanism of CSD initiation specific of neocortex, which is involved in the pathological mechanism of FHM3 mutations. Migraine etiology is multifactorial, with probably numerous different mechanisms (Ferrari et al., 2015); this novel mechanism of CSD initiation may be implicated also in other types of migraine, because it could be associated not only to Na_V_1.1 mutations, but also to other dysfunctions that lead to GABAergic neurons’ hyperactivity.

## Acknowledgements

We thank Frederic Brau for his skillful expertise on imaging and image processing, Michelle Studer and Pascal Ezan for advices on immunohistochemistry experiments, Etienne Audinat for insightful discussions, and Yuchio Yanagawa for donating GAD67-GFP knock-in mice.

This research was supported by the Investissements d’Avenir-Laboratory of Excellence “Ion Channels Science and Therapeutics” (grant LabEx ICST ANR-11-LABX-0015-01 to MM) and the European Union projects DESIRE (grant n. EFP7-602531 to MM). SZ was a Ph.D. student of the “Ecole doctorale 85” and received a fellowship from the French Research Ministry (MENRT). Our laboratory is member of the “Fédération Hospitalo-Universitaire” FHU-INOVPAIN.

## Author Contribution

Performed experiments and collected data: OC, SZ, PS, AL, MA, SC

Analyzed and interpreted the data: OC, SZ, PS, AL, FD, SC, MM

Implemented the computational model: LL, MK, MD, MM

Designed the experiments: OC, SZ, SC, MM

Prepared the figures: OC, SZ, PS, LL, AL, SC, MM

Wrote the manuscript: OC, MM

## Supplementary Material

**Supplementary Figure 1.**
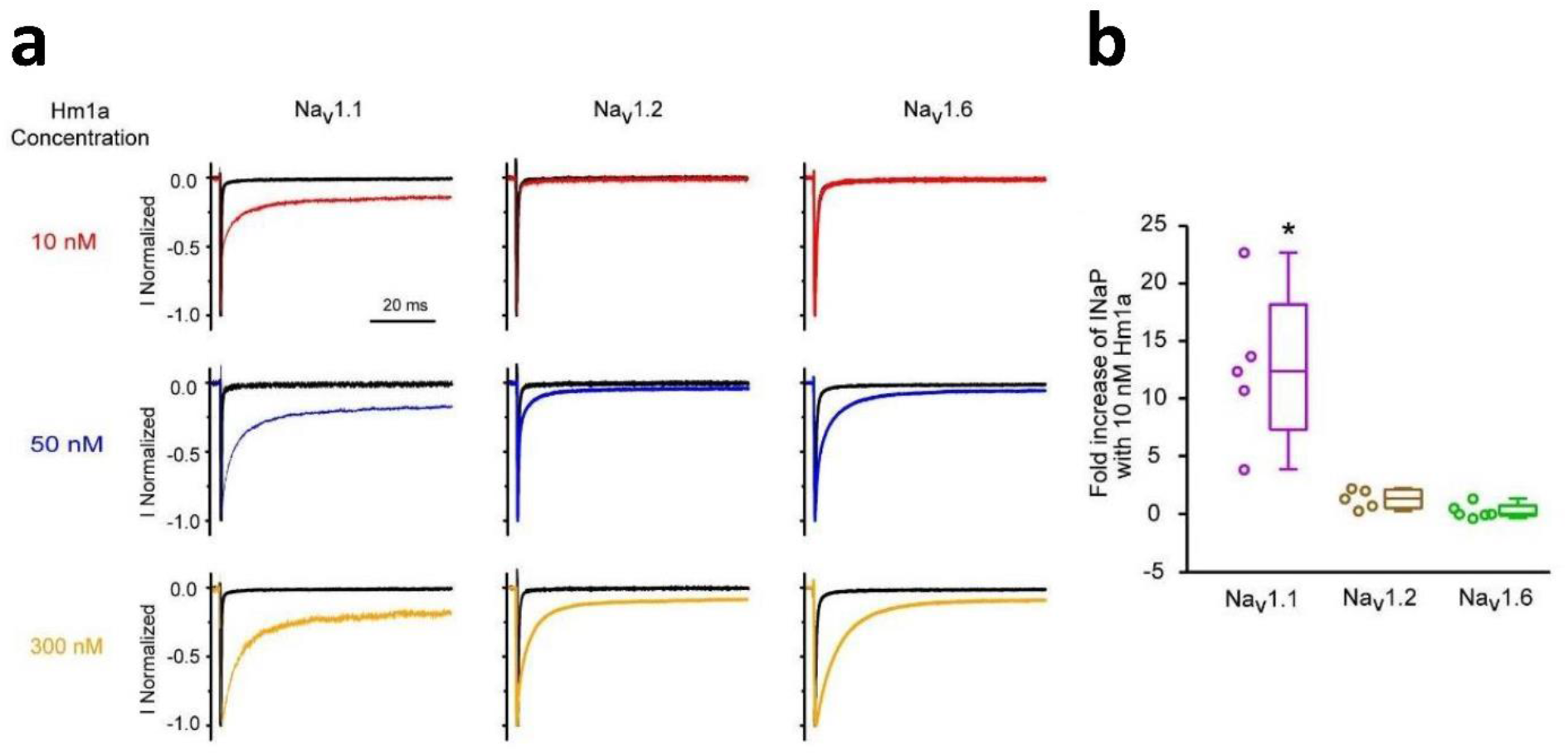
Test of the selectivity of our synthetic Hm1a peptide on Na^+^ channel isoforms expressed in the central nervous system: Na_v_1.1, Na_v_1.2 and Na_v_1.6. **a.** Representative whole-cell patch-clamp recordings of sodium currents evoked with 100ms depolarizing steps to −10mV from a holding potential of -100mV in tsA-201 cells expressing Na_v_1.1, Na_v_1.2, or Na_v_1.6 in control condition or in the presence of 10, 50 or 300nM Hm1a. Apparent EC50 (efficacy concentration at 50% of maximum effect) was 3nM for Na_v_1.1 (n=5 cells), 53nM for Na_v_1.2 (n=5 cells) and 41nM for Na_v_1.6 (n=6 cells). Note the selective enhancing effect of 10nM Hma1 on Na_v_1.1 compared to Na_v_1.2 and Na_v_1.6. At higher concentrations, Hma1 loses its selectivity on Na_v_1.1. **b.** Quantification of the effect of 10nM Hm1a displayed as fold increase of I_NaP_, showing that Hm1a is selective towards Na_v_1.1 over Na_v_1.2 and Na_v_1.6 isoforms. Whiskers box-chart plots represent median, minimum and maximum. One-sided one sample Wilcoxon Signed Rank Test, * p=0.029.

**Supplementary Figure 2.**
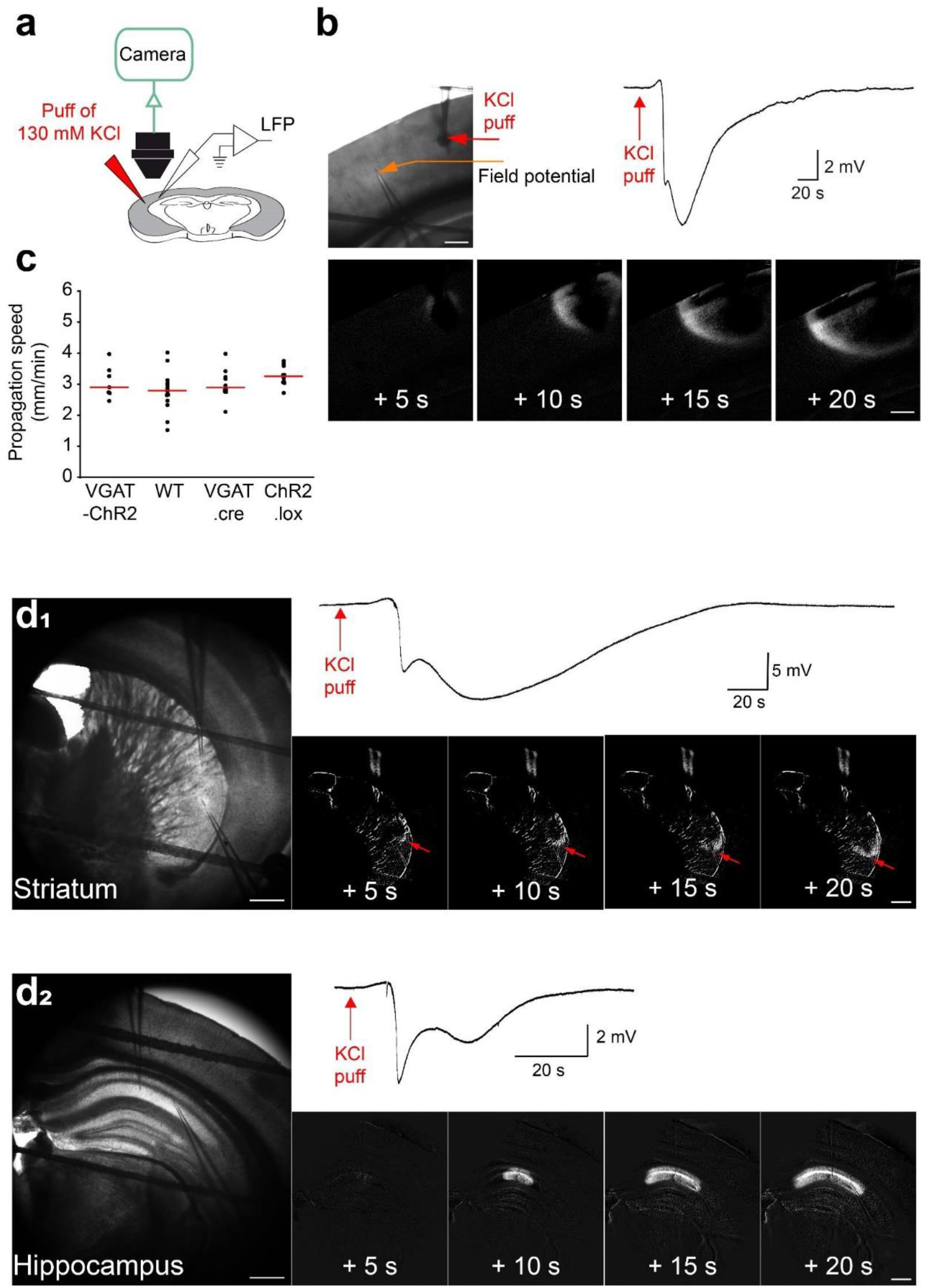
CSD is readily triggered by focal puffs of KCl 130mM in neocortex, striatum and hippocampus. **a**. Experimental design of KCl-induced CSD, which was triggered by a focal puff of 130 mM KCl applied using a picopump (10PSI, 500ms). Fastgreen (0.1%) was added to the KCl solution to visualize the effective KCl injection and quantify its area (mean=0.030 ± 0.005mm^2^, n=10 slices). **b.** Representative CSD induced in the neocortex and revealed by both a negative DC shift in the LFP and the propagating wave observed with intrinsic optical signal (IOS) imaging. The four lower panels show representative IOS images of CSD at different time points (image processing of raw images to better highlight the CSD wave, see methods). Scale bars 250 µm **c**. CSD was induced by 130mM KCl and had similar properties in different mouse lines: the dot plot displays the propagation speed in slices from VGAT-ChR2 mice (line that we used for optogenetic experiments, in which CSD has been induced with 130mM KCl after unsuccessful optogenetic illuminations; n=7 slices), WT littermate mice (n=14), VGAT.cre mice (n=10) and ChR2.lox mice (n=10) (Kruskall-Wallis test p=0.32); for the last three conditions the slices are those presented in Fig.3d (success rate plot). Bars correspond to medians. Pooling all the data, the CSD ignited by the KCl puff propagated in the cortical tissue at the speed of 3.01 ± 0.08mm/min (mean ± SEM, n=41 slices). **d_1_**. Representative CSD induced in the dorsal striatum by a puff of 130mM KCl as in b for the neocortex (n=7, 100% success rate); transmitted light image of the imaged area (left panel) and time series of processed images (right lower panels); the right upper panel shows the LFP recording; the CSD wave front is indicated by the red arrows. Scale bars 500µm **d_2_**. Representative CSD induced in the hippocampus by a puff of 130mM KCl as in b for the neocortex (n=8; 100% success rate); transmitted light image of the imaged area (left panel) and time series of processed images (right lower panels); the right upper panel shows the LFP recording. Scale bars: 500µm.

**Supplementary Figure 3.**
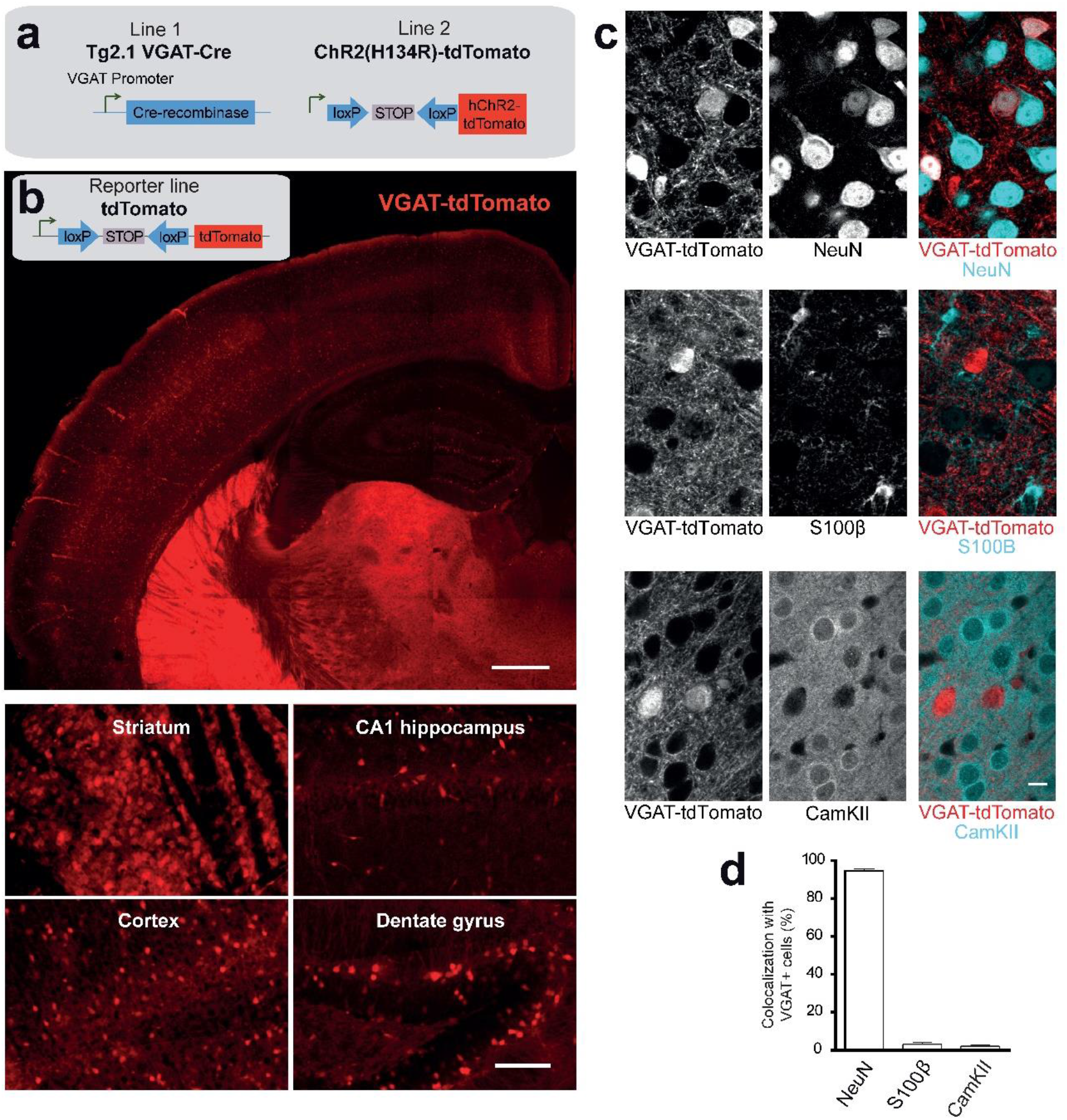
ChR2 is expressed selectively in GABAergic neurons and in numerous brain areas of VGAT-ChR2 mice. **a.** Mouse lines crossed for generating VGAT-ChR2 mice used for optogenetic experiments. **b.** Extended mosaic confocal image of a representative coronal brain slice from the mouse reporter line VGAT-tdTomato, in which VGAT-Cre+ cells (i.e. GABAergic neurons) express the cytoplasmatic tdTomato fluorescent protein (red), showing that the VGAT promoter drives expression in numerous brain areas including neocortex, hippocampus, striatum and the thalamus neuropil. Scale bar: 500µm. The lower panels display magnified areas from the upper panel, with single cell resolution. Scale bar: 100µm **c.** Representative confocal images of neocortical immunostainings with a neuronal marker (NeuN), an astrocytic marker (S100β) and a marker of excitatory neurons (CamKIIa) in tissue from VGAT-tdTomato mice. Scale bar: 10µm **d.** Quantifications of NeuN+ cells (95 ± 1%, grand average of 3 mice), S100β+ cells (3.0 ± 0.9%, grand average of 3 mice), and CamKII+ cells (2.0 ± 0.5%, grand average of 3 mice) among neocortical VGAT-dtTomato cells, confirming that ChR2 in VGAT-ChR2 mice is not expressed in excitatory neurons and astrocytes. We have not used VGAT-ChR2 mice in these experiments because the fluorescence of tdTomato-tagged ChR2 is mainly limited to the plasma membrane, and thus difficult to identify for an accurate quantification at the cellular level.

**Supplementary table.** Dynamics of CSD induced by optogenetic stimulations. **a.** Success rate (%), propagation speed (mm/min) and initiation latency (seconds) of CSD induced by 470 nm illumination in VGAT-ChR2 slices (Ctrl), VGAT-ChR2 *Scn1a* ^+/−^ slices in control conditions and VGAT-ChR2 *Scn1a*^+/−^ slices after [K^+^]_out_ increase to 8 mM. **b.** The same measurements in VGAT-ChR2 slices treated with blockers of the K^+^-Cl^−^ cotransporter KCC2 (VU024551-10 μM or VU0463271-10 μM) alone or with a GABA-A receptor agonist (Isoguvacine-10 μM), used to increase KCC2 baseline activity. Note that some of these results have been already presented in Fig.4. “n” refers to the number of slices, and “NA” to data not available (see methods).

|  | Ctrl | Na<sub>v</sub>1.1<sup>+/-</sup> | Na<sub>v</sub>1.1<sup>+/-</sup> + ACSF 8mM KCl |
| --- | --- | --- | --- |
| n | 27 | 12 | 7 |
| Mean | 2.41 | 3.20 | 3.97 |
| S.E.M | 0.15 | 0.20 | 0.28 |
| Median | 3.07 | 3.12 | 3.95 |
| NA | 0 | 0 | 0 |
| Statistics | Kruskal-wallis p=0.07 |  |  |

Latency of CSD induction (in seconds)
|  | Ctrl | Na<sub>v</sub>1.1<sup>+/-</sup> | Na<sub>v</sub>1.1<sup>+/-</sup> + ACSF 8mM KCl |
| --- | --- | --- | --- |
| n | 23 | 11 | 6 |
| Mean | 39 | 45 | 51 |
| S.E.M | 4 | 5 | 14 |
| Median | 39 | 38 | 47 |
| NA | 4 | 1 | 1 |
| Statistics | Kruskal-wallis p=0.70 |  |  |

NA = Data Not Available

**b** Success rate
|  | Ctrl | VU0240551 10μM | VU0463271 10μM | Isoguvacine 10μM | VU0463271 10μM + IsoG 10μM |
| --- | --- | --- | --- | --- | --- |
| n | 39/39 (100%) | 12/12 (100%) | 17/20 (85%) | 11/11 (100%) | 9/10 (90%) |
| Statistics | Fisher's exact test p=0.049 |  |  |  |  |
| Pairwise test adjusted with Bonferroni correction | NS | NS | NS | NS | NS |

Propagation speed of CSD (mm/min)
|  | Ctrl | VU0240551 10μM | VU0463271 10μM | Isoguvacine 10μM | VU0463271 10μM + IsoG 10μM |
| --- | --- | --- | --- | --- | --- |
| n | 38 | 12 | 15 | 11 | 9 |
| Mean | 3.38 | 3.12 | 3.32 | 3.08 | 3.19 |
| S.E.M | 0.13 | 0.26 | 0.18 | 0.25 | 0.16 |
| Median | 3.42 | 2.89 | 3.28 | 3.12 | 3.00 |
| NA | 1 | 0 | 2 | 0 | 0 |
| Statistics | Kruskal-wallis p=0.54 |  |  |  |  |

Latency of CSD induction (in seconds)
|  | Ctrl | VU0240551 10μM | VU0463271 10μM | Isoguvacine 10μM | VU0463271 10μM + IsoG 10μM |
| --- | --- | --- | --- | --- | --- |
| n | 37 | 12 | 11 | 11 | 9 |
| Mean | 20.0 | 21.0 | 16.5 | 19.0 | 24.0 |
| S.E.M | 2.3 | 4.2 | 3.5 | 3.5 | 9.1 |
| Median | 14.0 | 14.0 | 13.0 | 15.0 | 12.0 |
| NA | 2 | 0 | 6 | 0 | 0 |
| Statistics | Kruskal-wallis p=0.93 |  |  |  |  |

